# Structured navigation emerges from self-guided spatial learning in freely moving common marmosets

**DOI:** 10.1101/2025.03.22.644580

**Authors:** Nada El Mahmoudi, Francesca Lanzarini, Farzad Ziaie Nezhad, Deepak Surendran, Jean Laurens

**Affiliations:** Ernst Strüngmann Institute (ESI) for Neuroscience in Cooperation with Max Planck Society, Frankfurt am Main, 60528, Germany

**Author notes:** Co-first authors.

**Keywords:** Common marmosets, Naturalistic Behavior, Spatial memory, Navigation strategies, Self-guided learning, Topological organization, Behavioral structure, Non-Human Primates, Neuroethology

## Abstract

How do structured, memory-guided behaviors emerge in freely moving primates? We addressed this question by training common marmosets to perform a foraging task in a semi-naturalistic environment, where they retrieved food from eight dispensers over multiple sessions. Animals received no instruction or shaping and were free to visit dispensers in any order. We found that animals engaged with the task in most trials and, within engaged trials, transitioned from exploratory to efficient foraging behavior. Learning was marked by performance gains and the emergence of reproducible movement patterns between specific dispensers. Using trajectory analyses and probabilistic modeling, we found that animals formed stable route segments linking specific dispensers, reflecting the emergence of local navigation motifs. These segments became increasingly regular and predictable with experience. Yet rather than being rigidly replayed, they were flexibly recombined into variable global sequences. This indicates that animals adopted a hybrid navigation strategy, in which reusable route segments are embedded within a topological structure. These findings demonstrate how efficient, adaptive navigation can emerge through self-guided experience in complex environments. Our approach provides a naturalistic and longitudinal framework for studying the formation of structured spatial strategies in non-human primates, bridging ecological behavior with theoretical models of learning and memory.

## Introduction

How structured behavior emerges from repeated experience in complex environments is a fundamental question in cognitive neuroscience and behavioral ecology. Navigation provides a powerful model to address this question, as it requires animals to extract, organize, and flexibly use spatial information across multiple timescales and contexts.

Although the cognitive and neural foundations of navigation have been extensively studied in rodents, how these mechanisms operate in non-human primates (NHPs) remains poorly understood. Primates differ markedly from rodents in their sensorimotor capacities, brain architectures, and ecological demands ^1–5^, differences that become especially relevant when studied under ethologically valid conditions.

Understanding how spatial memory supports navigation in primates, and whether similar organizational principles apply across species, is a key step toward bridging this gap. Spatial memory enables animals to remember where they have been, and to use that information to guide future movement ^6^. It is classically divided into spatial working memory, supporting short-term retention during ongoing behavior, and spatial reference memory, which stores stable knowledge across repeated experience ^7^.

These memory systems support a range of navigation strategies, defined by how spatial information is structured and used to generate behavior ^8–10^. Route-based navigation relies on learned sequences of subgoals; topological navigation involves representations of interconnected spatial nodes; and metric navigation supports flexible rerouting based on distances and directions ^11,12^.

NHPs display remarkable spatial flexibility and foraging efficiency ^4^, yet the mechanisms by which such navigation strategies emerge and evolve over time remain elusive. Experimental approaches to primate navigation have traditionally relied on simplified laboratory tasks, which limit ecological validity, or on observational field studies, which preclude high-resolution behavioral tracking. As a result, we lack a detailed understanding of how spatial learning unfolds in freely moving primates.

The common marmoset (*Callithrix jacchus*) provides a powerful model to address this gap. These arboreal primates navigate three-dimensional environments to access both stable and unpredictable food sources, such as exudates, fruits, and insects ^13–17^. In the wild, they follow repeatedly used foraging routes embedded within topological maps, enabling flexible yet structured navigation ^18^. However, how such strategies are acquired and optimized over time, and whether similar processes can be observed under controlled yet ethologically relevant conditions, remains unclear.

To address this, we developed a semi-naturalistic foraging task in which marmosets freely visited eight spatially distributed food dispensers to collect rewards, without external instruction or shaping. By combining behavioral clustering, trajectory analysis, and probabilistic modeling, we showed how freely moving marmosets progressively structured their navigation behavior through repeated experience. With training, animals developed increasingly efficient navigation patterns, characterized by the formation and repeated use of stable route segments linking specific dispenser locations. However, rather than rigidly replaying fixed routes, they flexibly recombined these segments into variable global trajectories, revealing a balance between behavioral consistency and strategic flexibility consistent with topological navigation. Our results also show that behavioral variability in late training reflected a shift from generating novel sequences to reusing familiar ones, suggesting a transition from exploration to flexible exploitation.

Altogether, our findings provide a proof of concept for using naturalistic behavioral paradigms to uncover the emergence and consolidation of memory-guided navigation strategies in non-human primates. Our framework bridges ecological behavior and theoretical models of spatial cognition, offering new insight into the cognitive foundations of primate navigation.

## Results

We implemented a spatial memory task in an enriched, semi-naturalistic environment to study navigation behavior in common marmosets (**Fig. 1A**). Animals moved freely within the arena and collected rewards from eight food dispensers arranged around its perimeter (**Fig. 1E**). Each trial began when the central “Home Dispenser” was activated, signaled by auditory and visual cues along with reward delivery. Upon visiting this dispenser, all eight peripheral dispensers became simultaneously active and dispensed reward (**Fig. 1B**). A trial was completed when the animal had visited all eight unique dispensers, regardless of any revisits along the way. If this condition was not met within 10 minutes, the trial ended automatically. The next trial began with the reactivation of the Home Dispenser. Each subject engaged with the task over 30 sessions, performing on average 7.96 ± 0.4 trials per session for Animal E and 8 ± 0.4 for Animal F.

**Figure 1:**
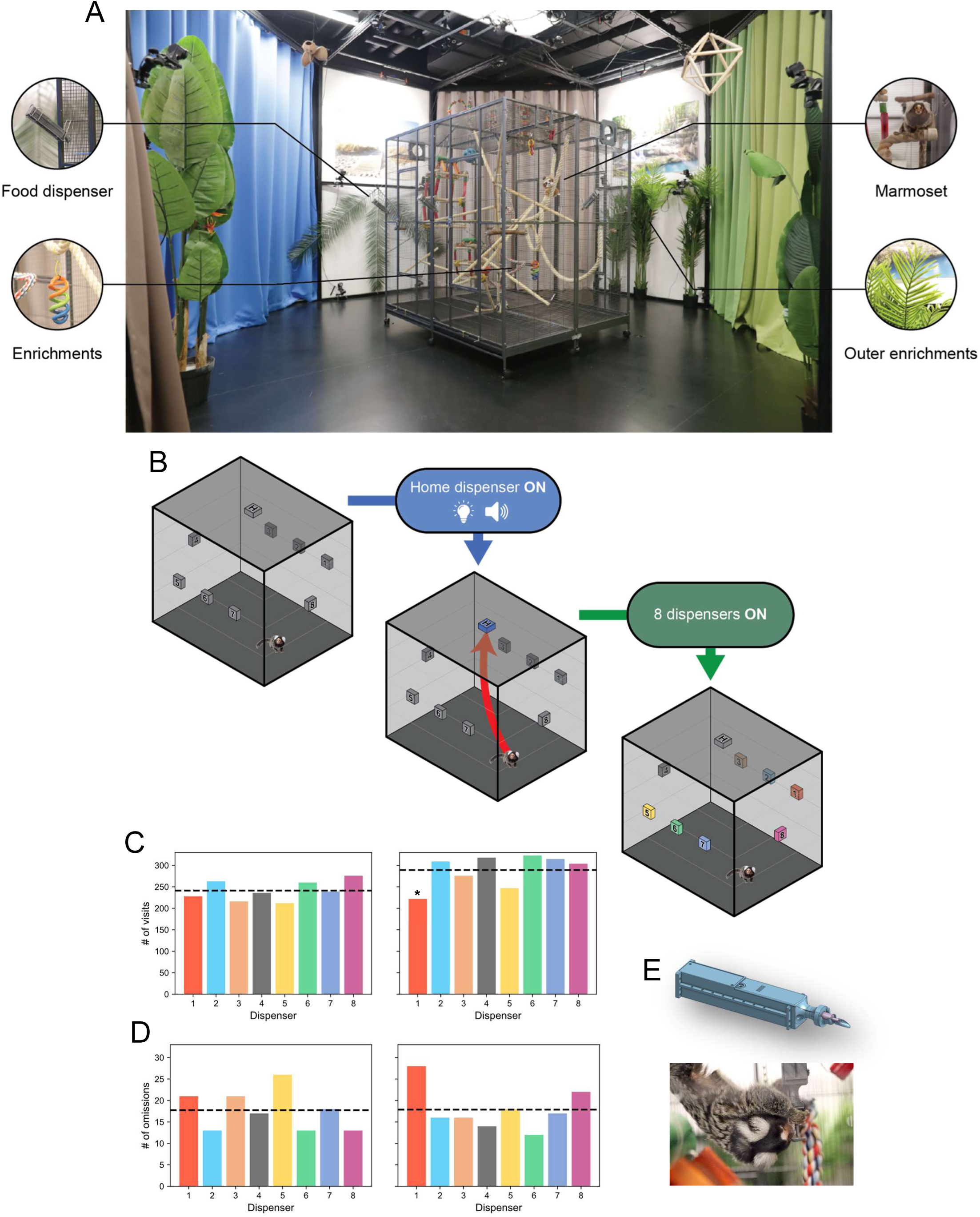
A freely moving setup for investigating spatial memory in marmosets. **A. Experimental setup**. The enclosure consisted of a wire mesh cage (1.8 m × 1.4 m × 1.6 m; 4 m³) that allowed the marmosets to climb and move freely. Structural enrichments (wooden sticks, semi-rigid ropes, and various objects) were included to promote locomotion and exploration. External spatial cues were provided by four white walls decorated with distinct landscape posters, four colored curtains, and large plastic plants. An action camera was mounted above the setup to record behavior. A centrally mounted food dispenser, designated the Home Dispenser (HD), was placed on the ceiling, while eight peripheral food dispensers were evenly distributed around the perimeter of the cage to deliver rewards during the task. **B. Trial design.** At the start of each trial, the HD was activated with an LED and a buzzer. Once the marmoset reached the HD, all eight peripheral dispensers were simultaneously activated and delivered gum arabic. To complete the trial, the animal had to visit each of the eight dispensers at least once. If not completed within 10 minutes, the trial was automatically terminated. **C. Total visits per dispenser.** Bar plots showing the total number of visits to each of the eight dispensers for Animal E (*left*) and Animal F (*right*). The dashed horizontal line indicates the expected number of visits under a uniform distribution. A chi-squared test assessed the overall distribution, followed by binomial tests comparing each dispenser to the expected proportion. Significant p-values (after Bonferroni correction) are indicated with an asterisk (*p < 0.05). **D. Total omissions per dispenser.** Bar plots showing the total number of omissions for each dispenser in Animal E (*left*) and Animal F (*right*). The dashed horizontal line indicates the expected number of omissions under a uniform distribution. A chi-squared test was used to assess deviation from uniformity. **E. Reward dispenser design.** Design of the custom-made food dispenser used during the experiment (*top*). Eight dispensers were placed around the setup, while the HD was placed at the center of the ceiling upside-down (*bottom*, a marmoset eating from the HD).

### Marmosets show stable dispenser interactions and distinct learning dynamics across error types

To assess marmoset interactions with the dispensers, we analyzed the total number of visits and omissions (i.e., dispensers not visited within a trial) for each dispenser (**Fig**. **1C-D**). A Chi-squared test revealed significant deviations from uniformity in the distribution of visits for both Animal E (χ^2^ = 15.47, df = 7, p = 0.03) and Animal F (χ^2^ = 33.6, df = 7, p < 0.001). Post hoc binomial tests with Bonferroni correction, however, indicated that Animal E visited all dispensers at comparable rates (p > 0.05), while Animal F visited Dispenser 1 significantly less often than expected (p < 0.05), with no significant differences among the remaining dispensers. For omissions, neither animal showed significant deviations from uniformity (Animal E: χ^2^ = 8.9, df = 7, p = 0.26, Animal F: χ^2^ = 9.89, df = 7, p = 0.19), suggesting no systematic neglect of specific locations. Together, these results indicate that dispenser interactions were globally stable and well-balanced across the arena, providing a robust basis for analyzing how spatial behavior and error patterns evolve over time.

To evaluate task performance, we first classified the errors observed during the trials into three types: *repetition errors* (revisiting a dispenser already visited during the same trial), *omission errors* (failing to visit all eight dispensers), and *rule errors* (returning to the home dispenser mid-trial). We then computed the mean number of total errors per session and assessed how this measure evolved over time using linear regression (**Fig. S1A**). Both animals exhibited a significant decrease in error rate across sessions, reflecting a spontaneous improvement in performance (Animal E: R^2^ = 0.26, p < 0.01, β = -0.07, Animal F: R^2^ = 0.62, p < 0.001, β = -0.28). To assess potential contributors to this improvement, we used a multiple linear regression model (**Table S1**), which confirmed that the number of sessions was the main predictor of performance gains, with no detectable contribution from other variables. To further describe the evolution of different error types over time, we compared their session-wise dynamics using a Kruskal-Wallis test (**Fig. S1B**). This analysis revealed that the temporal profiles of repetition, omission, and rule errors differed significantly (Animal E: H = 13.06, *p* < 0.01, Animal F: H = 39.05, *p* < 0.001), suggesting that the different error types did not follow the same trajectory across training. Finally, to examine how error types related to each other and to overall performance, we conducted a correlation matrix analysis (**Fig. S1C-D**). Repetition and rule errors were positively correlated with each other and with the total number of visits, suggesting they tended to co-occur in more exploratory or disorganized trials. In contrast, omission errors were uncorrelated with both repetition and rule errors, suggesting that they may reflect distinct behavioral processes.

### Clustering analysis reveals different task execution profiles

Based on this dissociation, we applied an unsupervised k-means clustering analysis to capture trial-by-trial variability in task execution. We used omission errors and total visit counts as two complementary metrics, indexing disengagement and activity level, respectively. Although moderately correlated (**Fig. S1C-D**), these measures captured distinct behavioral dimensions, enabling the identification of discrete behavioral states.

The optimal number of clusters (*k = 4*) was determined using silhouette score and inertia, which quantify within-cluster coherence and between-cluster separation (**Fig. 2A**). For both animals, clustering produced coherent and interpretable partitions (Animal E: silhouette = 0.73, inertia = 41.9, Animal F: silhouette = 0.68, inertia = 56.4), reflecting consistent behavioral variability across trials. Cluster centroids are reported in **Table S2**. To interpret the behavioral significance of each cluster, we compared key performance metrics across them (**Fig. S3**):

- **Cluster 0** exhibited the fewest errors and shortest trial durations (p < 0.001 for both animals vs. all other clusters), reflecting a state of optimal performance marked by efficient and precise behavior.
- **Cluster 1** was associated with higher visit counts and increased rule and repetition errors (p < 0.001), but few omissions, suggesting active but imprecise task engagement.
- **Cluster 2** showed the highest omission rates and no dispenser visits, consistent with disengagement or lack of task initiation.
- **Cluster 3** reflected partial engagement, with fewer visits and more omissions than Clusters 0 and 1 (p < 0.001), but slightly more visits and fewer omissions than Cluster 2 (non-significant), indicating incomplete task execution.

**Figure 2:**
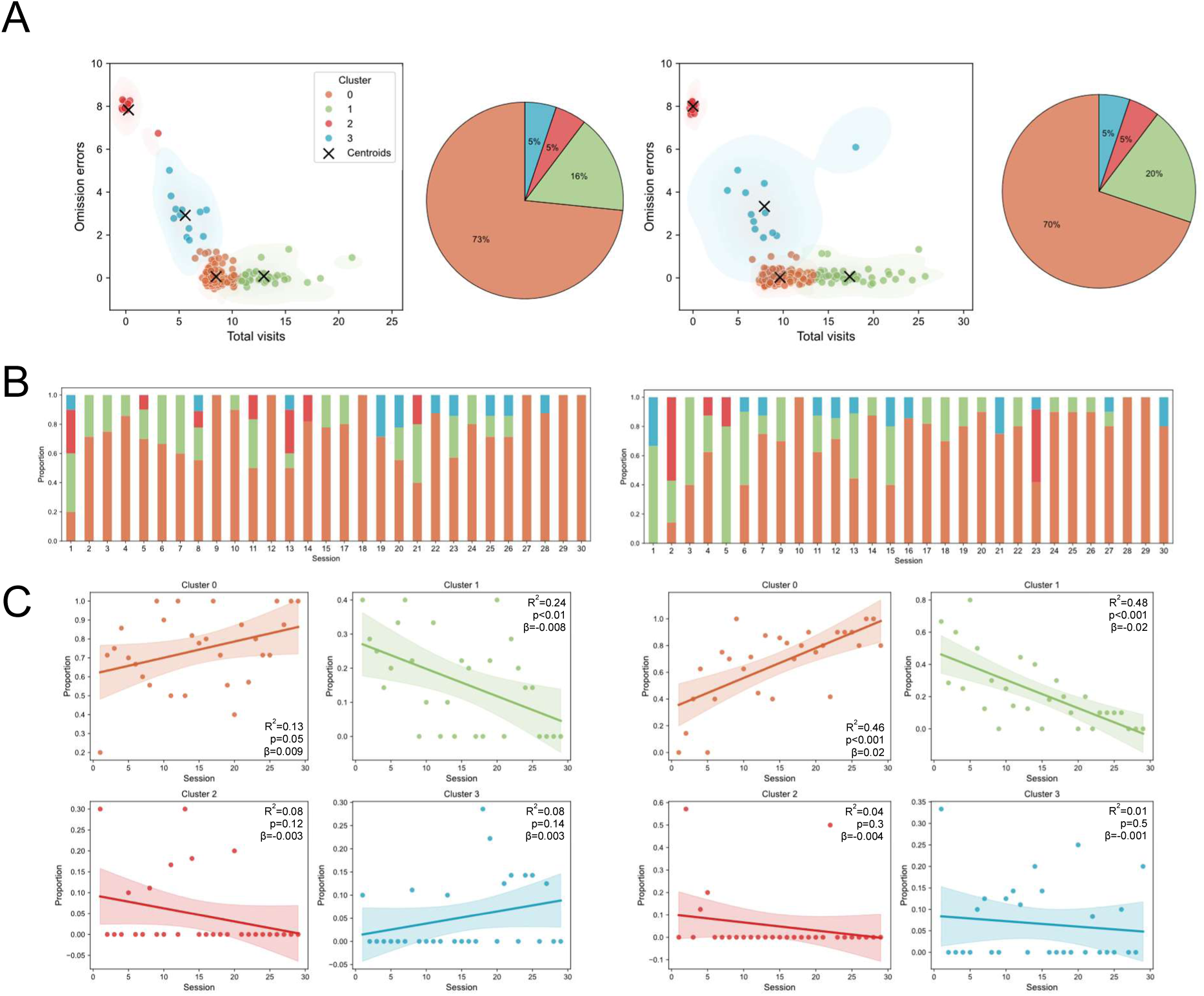
Clustering analysis reveals different task execution profiles. **A. Unsupervised clustering analysis.** Scatter plots showing the outcome of the k-means clustering analysis for Animal E (*left*) and Animal F (*right*), with each trial assigned to one of four clusters (color-coded). Cluster centroids, defined by omission errors and total number of visits, are marked with crosses. Data points are jittered for visibility. Pie charts indicate the proportion of trials assigned to each cluster. Colors are consistent across panels: Cluster 0 (orange), Cluster 1 (green), Cluster 2 (red), and Cluster 3 (blue). **B. Cluster distributions across sessions.** Stacked bar plots showing the normalized proportion of trials in each cluster across training sessions for Animal E (*left*) and Animal F (*right*). Each bar represents one session, and segment height indicates the proportion of trials falling into each cluster. **C. Temporal evolution of cluster proportions**. Scatter plots showing the proportion of trials in each cluster across sessions. Dashed lines represent regression fits; shaded areas indicate 95% confidence intervals. R^2^, p-values, and regression slope (β) quantify the strength and direction of the association.

Cluster distribution was remarkably consistent between animals: Clusters 0 and 1 together accounted for the vast majority of trials (89% for Animal E, 90% for Animal F), while Clusters 2 and 3 comprised only 10% each, suggesting that most trials reflected engaged task performance (**Fig. 2A**).

We next assessed how cluster distributions evolved over time (**Fig. 2B-C**). Linear regressions showed that the proportion of Cluster 0 trials increased with session number (Animal E: R^2^ = 0.13, p = 0.05, Animal F: R^2^ = 0.45, p < 0.001), while Cluster 1 decreased (Animal E: R^2^ = 0.24, p < 0.01, Animal F: R^2^ = 0.48, p < 0.001), reflecting a gradual shift from error-prone to optimized task execution. In contrast, the frequency of Clusters 2 and 3 remained stable over time (p > 0.1), suggesting that disengagement levels were not affected by session progression.

Altogether, these findings support a functional distinction between clusters reflecting task-related engagement (Clusters 0 and 1) and those associated with limited or absent task participation (Clusters 2 and 3), enabling the identification of distinct behavioral states during task performance.

### Clusters 0 and 1 reflect variations in efficiency within a shared behavioral framework

Since Clusters 0 and 1 were associated with task engagement, we next asked whether they reflected distinct navigation strategies or simply differences in execution efficiency. We focused on the first eight choices within each trial, as successful task completion required visiting eight unique dispensers without repetition.

We quantified the performance rate as the percentage of correct responses within the first eight choices of each trial (**Fig. 3A, H**), where a response was considered correct if the animal visited a dispenser that had not yet been visited during that trial. To determine whether performance deviated from random behavior, we compared performance rates to an expected chance level of 50%, based on uniform random choice among all dispensers at each step (see Methods). Wilcoxon signed-rank test with Bonferroni correction confirmed that both clusters exhibited significantly greater performance rates than chance (p < 0.001, for both animals), confirming that even in Cluster 1, despite lower precision, marmosets showed goal-directed behavior.

**Figure 3:**
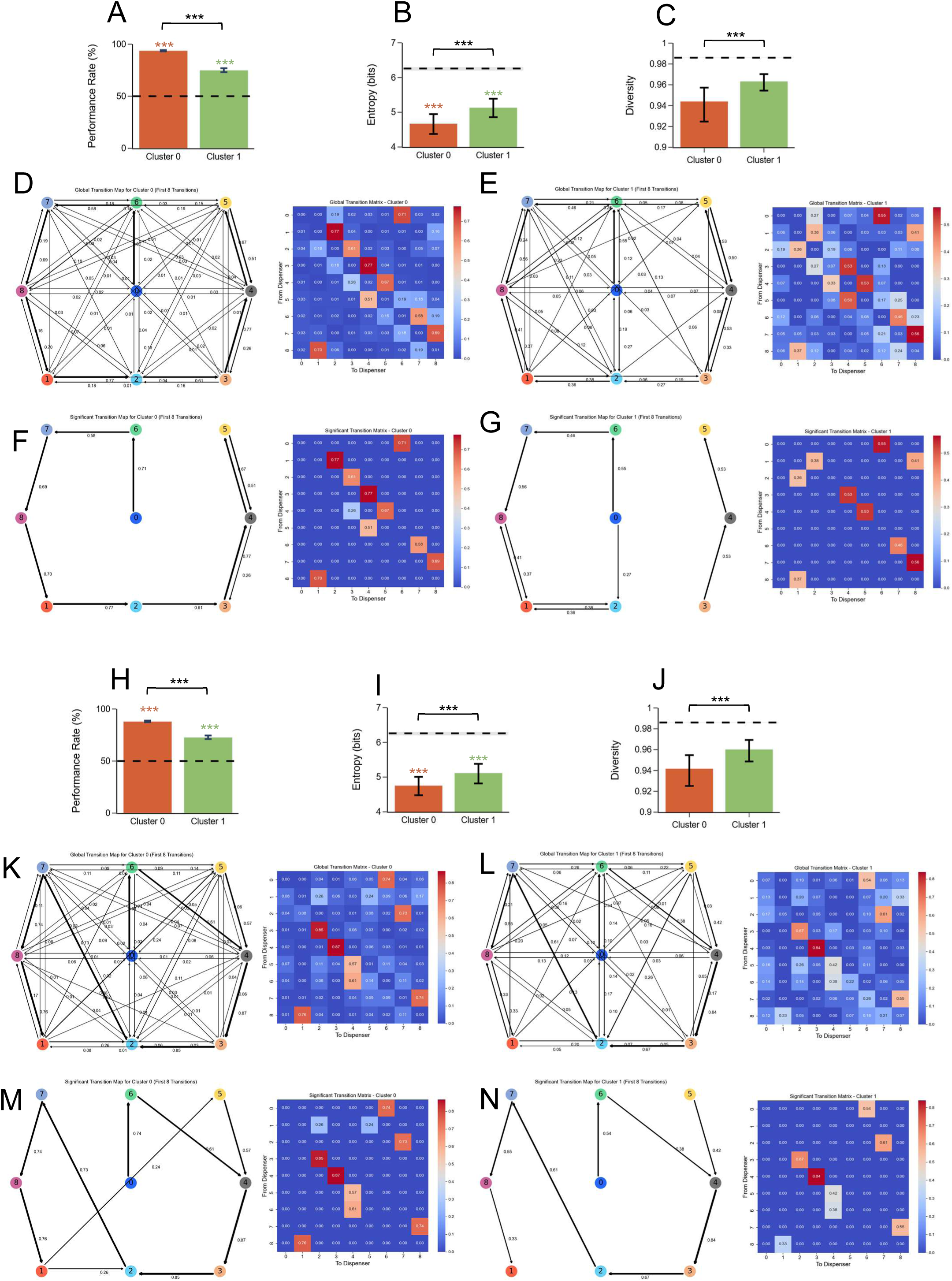
Clusters 0 and 1 reflect variations in efficiency within a shared behavioral framework. *(A–G) Analyses for Animal E; (H–N) Corresponding analyses for Animal F.* A and H. Mean performance in the first eight choices for Clusters 0 and 1. Bar plots show the percentage of correct responses (i.e., visits to non-repeated dispensers) within the first eight choices of each trial. Error bars indicate SEM. The dashed line marks the 50% chance level (random choice among nine options). Wilcoxon signed-rank tests (vs. chance) and Mann-Whitney U tests (between clusters) were performed, with Bonferroni correction. ***p < 0.001. **B and I. Global entropy across Clusters 0 and 1.** Bar plots display mean global entropy (in bits), with 95% bootstrap confidence intervals (CI). Dashed lines represent mean entropy expected under random transitions; shaded areas indicate 95% CI of random baseline. Permutation tests with Bonferroni correction assessed significance. ***p < 0.001. **C and J. Global transition diversity across Clusters 0 and 1**. Same conventions as in B and I. Bars represent mean transition diversity; dashed lines and shaded areas reflect the random baseline. ***p < 0.001. **D and K. Global transition maps and matrices for Cluster 0**. *Left:* Directed graphs represent all observed transitions within the first eight choices. Node positions correspond to dispenser locations and arrow thickness is proportional to transition probability. *Right*: transition matrices show the probability of moving between all dispenser pairs, with color intensity reflecting transition strength (dark red: high, dark blue: low). **E and L. Global transition maps and matrices for Cluster 1**. Same conventions as in D and K. **F and M. Statistically significant transition maps and matrices for Cluster 0**. *Left:* Directed graphs show transitions that occurred significantly more often than expected under a uniform random model (p = 1/9; binomial test with Bonferroni correction). *Right:* corresponding transition matrices highlight these statistically significant transitions. Arrow thickness and color intensity reflect transition frequency, with thicker arrows and warmer colors indicating higher occurrence rates. **G and N. Statistically significant transition maps and matrices for Cluster 1.** Same conventions as in F and M.

To further explore how they navigated the environment, we analyzed their transitions between dispensers within a trial, i.e., how they moved from one dispenser to another. By reconstructing sequences of dispenser visits, we captured the order in which animals visited dispensers. We quantified the global organization of transitions using two complementary measures: global entropy, capturing the uncertainty in transition patterns and transition diversity, reflecting the evenness of transition use. In our framework, high entropy reflects unpredictable, wide-ranging transitions, while low entropy indicates a reliance on a smaller set of recurrent transitions. Similarly, high diversity corresponds to evenly distributed transitions, whereas low diversity signifies a preference for specific paths. For both animals, entropy and transition diversity were significantly lower than chance (p < 0.001, bootstrap with 1,000 random transitions, **Fig. 3B-C, I-J)**, indicating structured and non-random transitions in both clusters.

To visualize these patterns, we built transition matrices and transition maps. Global transition maps showed all transitions, with arrow thickness indicating transition probabilities (**Fig. 3D-E, K-L**). Significant transition maps (based on binomial tests with Bonferroni correction) highlighted transitions occurring more frequently than expected by chance (binomial tests with Bonferroni correction, **Fig. 3F-G, 3M-N**). Correlation analyses revealed a strong alignment in transition structure between clusters (global matrices: Animal E: r=0.89 p<0.001, Animal F: r=0.91 p<0.001, significant matrices: Animal E: r=0.8 p<0.001, Animal F: r=0.96 p<0.001). Yet, Euclidean distances indicated differences in absolute transition probabilities (global matrices: Animal E: 0.90, Animal F: 0.84, significant matrices: Animal E: 1.92, Animal F: 0.72), suggesting that while the structure of transitions was conserved, the specific weights assigned to each path varied between clusters.

Altogether, these findings indicate that Clusters 0 and 1 share a common navigational structure, differing primarily in execution efficiency rather than in strategy type. Based on this similarity, we pooled both clusters for subsequent analyses of spatial learning.

### Evolution of performance and global transition pattern across training phases

We next examined whether improvements in task performance were accompanied by changes in the global organization of navigation behavior. To do so, we divided the 30 training sessions into six consecutive phases of five sessions each: early training (Beg 1: sessions 1-5, Beg 2: sessions 6-10), intermediate training (Int 1: sessions 11-15, Int 2: sessions 16-20), and late training (Last 1: sessions 21-25, Last 2: sessions 26-30).

As expected, performance, quantified as the percentage of correct responses within the first eight choices, significantly increased across training phases (**Fig. 4A, F**, Kruskal-Wallis test, Animal E: H = 11.66, p = 0.03, Animal F: H = 64.6, p < 0.001).

**Figure 4:**
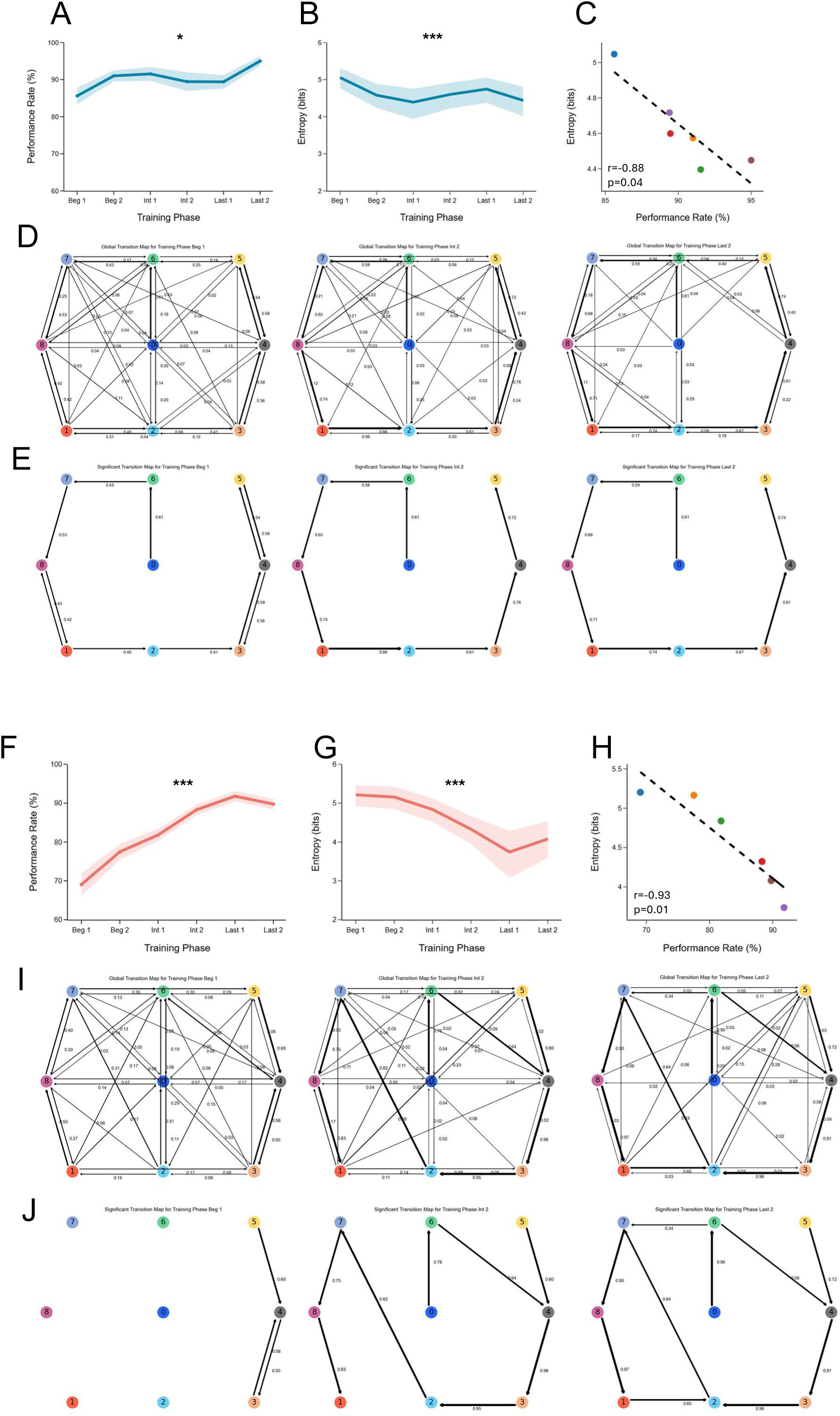
Evolution of performance and transition pattern across training phases. *(A–E) Analyses for Animal E; (F–J) Corresponding analyses for Animal F.* **A and F. Performance across training phases.** Line plots showing the mean performance (% correct choices within the first 8 visits per trial) across six training phases. Solid lines represent the mean; shaded areas indicate the standard error of the mean (SEM). Kruskal-Wallis test followed by Dunn’s post hoc test with Bonferroni correction was used for statistical comparisons. Significant results are marked: ***p < 0.001, **p < 0.01, *p < 0.05. **B and G. Global entropy across training phases.** Line plots showing the evolution of mean global entropy (in bits) over training. Points represent phase means; shaded areas represent 95% confidence intervals. Statistical analysis used permutation tests with Bonferroni correction. Asterisks indicate significant phase-wise differences. **C and H. Correlation between performance and entropy.** Scatter plots showing the relationship between performance rate and global entropy across phases. Each point represents a training phase, color-coded accordingly. Dashed black lines indicate linear regression fits. Pearson correlation coefficients are reported, with Bonferroni correction for multiple comparisons. **D and I. Global transition maps.** Directed graphs showing transition patterns across training phases (Beg 1: sessions 1–5; Int 2: sessions 16–20; Last 2: sessions 26–30). Node positions match dispenser locations; arrow thickness represents transition probabilities. Thicker arrows indicate more frequently used transitions. **E and J. Statistically significant transition maps.** Directed graphs showing transitions that occurred significantly more often than expected by chance (uniform model, p = 1/9). Significance assessed via binomial tests with Bonferroni correction. Maps are shown for Beg 1, Int 2, and Last 2. Thicker arrows highlight dominant, structured transitions.

In parallel, global transition entropy within the first eight choices significantly decreased over training (**Fig. 4B, G**, permutation test, p < 0.001 for all comparisons), indicating a progressive emergence of more structured behavioral patterns. Moreover, this reduction in entropy was strongly associated with improved performance (**Fig. 4C, H**): Pearson correlation analyses revealed a robust negative relationship between mean global entropy and performance across training phases (Animal E: r = -0.88, p = 0.04, Animal F: r = -0.93, p = 0.01).

To further characterize this evolution, we calculated the overlap index, a measure of similarity between entropy distributions across training phases (0 = no overlap, 1 = complete overlap, **Fig. S4A, F**). Early phases exhibited low overlap (Beg 1 vs. Int 2: Animal E = 0.15, Animal F = 0.004), reflecting a marked reorganization of transition patterns early in training. In contrast, overlap increased between later phases (Int 2 vs. Last 2: Animal E = 0.7, Animal F = 0.5), suggesting stabilization and refinement of navigation strategies over time.

Global (**Fig. 4D, I**) and significant (**Fig. 4E, J**) transition maps further illustrated this refinement. Permutation-based correlation analyses (Bonferroni-corrected) confirmed that the overall structure of transition matrices remained consistent across training phases (**Fig. S4B, G**). However, Euclidean distances between matrices revealed substantial changes in individual transition probabilities, which progressively decreased with training, indicating reduced variability in transition patterns.

Altogether, these results show that performance improvements were accompanied by a gradual reduction in transition entropy and variability, reflecting the emergence and consolidation of consistent, structured navigation patterns across training.

### Local transition analysis uncovers dispenser-specific variability within a globally structured transition behavior

To determine whether the global structuring of transitions also emerged at finer spatial scales, we examined local entropy (e.g. uncertainty in the pattern of transitions leaving a given dispenser) patterns at the level of individual dispensers (**Fig. S4C, H**). For both animals, local entropy significantly decreased over training (Kruskal-Wallis, Animal E: H = 12.6, p = 0.02, Animal F: H = 25.67, p < 0.001), indicating increasing transition regularity at the dispenser level. Post hoc comparisons confirmed significant reductions between early (Beg 1) and late (Last 2) training phases (Animal E: p = 0.04, Animal F: p = 0.04).

To evaluate the behavioral relevance of this local structuring, we computed the probability of making an error after visiting each dispenser (**Fig. S4D, I**). For both animals, error probability decreased significantly between early (Beg 1) and late (Last 2) training phases (Kruskal-Wallis, Animal E: H = 6.5, p = 0.01, Animal F: H = 7.6, p < 0.01). Furthermore, we found a strong positive correlation between mean local entropy and mean error probability across phases (Animal E: r = 0.90, p < 0.01, Animal F: r = 1.0, p < 0.001, **Fig. S4E, J**), indicating that lower local entropy was closely associated with improved performance.

We next assessed spatial heterogeneity in local structuring by comparing entropy values across dispensers to a baseline generated from 1,000 bootstrap-sampled random transitions (mean = 2.65 bits, 95% CI: 2.15-2.98). All dispensers showed entropy values below this baseline (**Fig S4C,H**), confirming structured, non-random transitions. However, the degree of structuring varied across dispensers, as reflected by the standard deviation of local entropy values (Animal E: 0.29 bits, Animal F: 0.35 bits), indicating spatial heterogeneity in transition predictability.

Altogether, these results show that transition structuring develops not only globally but also at finer spatial scales. While overall entropy declines with training, individual dispensers vary in their contribution to this structuring, suggesting that some locations support more stereotyped transitions, whereas others retain greater flexibility.

### Marmoset transition patterns show local predictability but global flexibility

We next asked whether the increasing structuring of transition patterns over time reflected the emergence of rigid, repetitive strategies, or instead more flexible, probabilistic preferences. Specifically, we examined whether successful performance in error-free trials consistently relied on the same transition patterns, or whether multiple strategies could lead to success, suggesting variability in execution despite overall structure.

To test this, we applied a first-order Markov chain model to assess the predictability of transitions at both local and global levels. Markov models estimate the probability of moving from one state (dispenser) to another based solely on the current state, and are well suited to capture sequential dependencies. This allowed us to evaluate the model’s ability to predict individual transitions (local level) as well as full sequences of dispenser visits (global level). The model was trained exclusively on error-free trials to isolate structured behaviors associated with successful task execution, and its performance was compared to a random baseline model assuming uniform transition probabilities.

The Markov model showed strong performance at the local level (**Fig. 5A-D**), with average prediction accuracy of 79.6% ± 7.8% (95% CI) for Animal E and 84.75% ± 2.7% for Animal F, well above the random baseline (Animal E: 10.5% ± 1.8%, Animal F: 9.0% ± 3.5%). Wilcoxon signed-rank tests confirmed that these differences were statistically significant (p < 0.01), indicating that local transitions were highly structured and predictable. To assess global predictability, we used the model to generate full action sequences by iteratively selecting the most probable next transition, starting from the first dispenser visited in each trial (**Fig. 5E-F**). As expected, accuracy declined relative to local prediction, but remained substantially above chance. Average full-sequence accuracy reached 57.1% ± 13.8% (Animal E) and 54.5% ± 6.5% (Animal F), while the random baseline achieved 0% for both animals. These differences were also statistically significant (p < 0.01).

**Figure 5.**
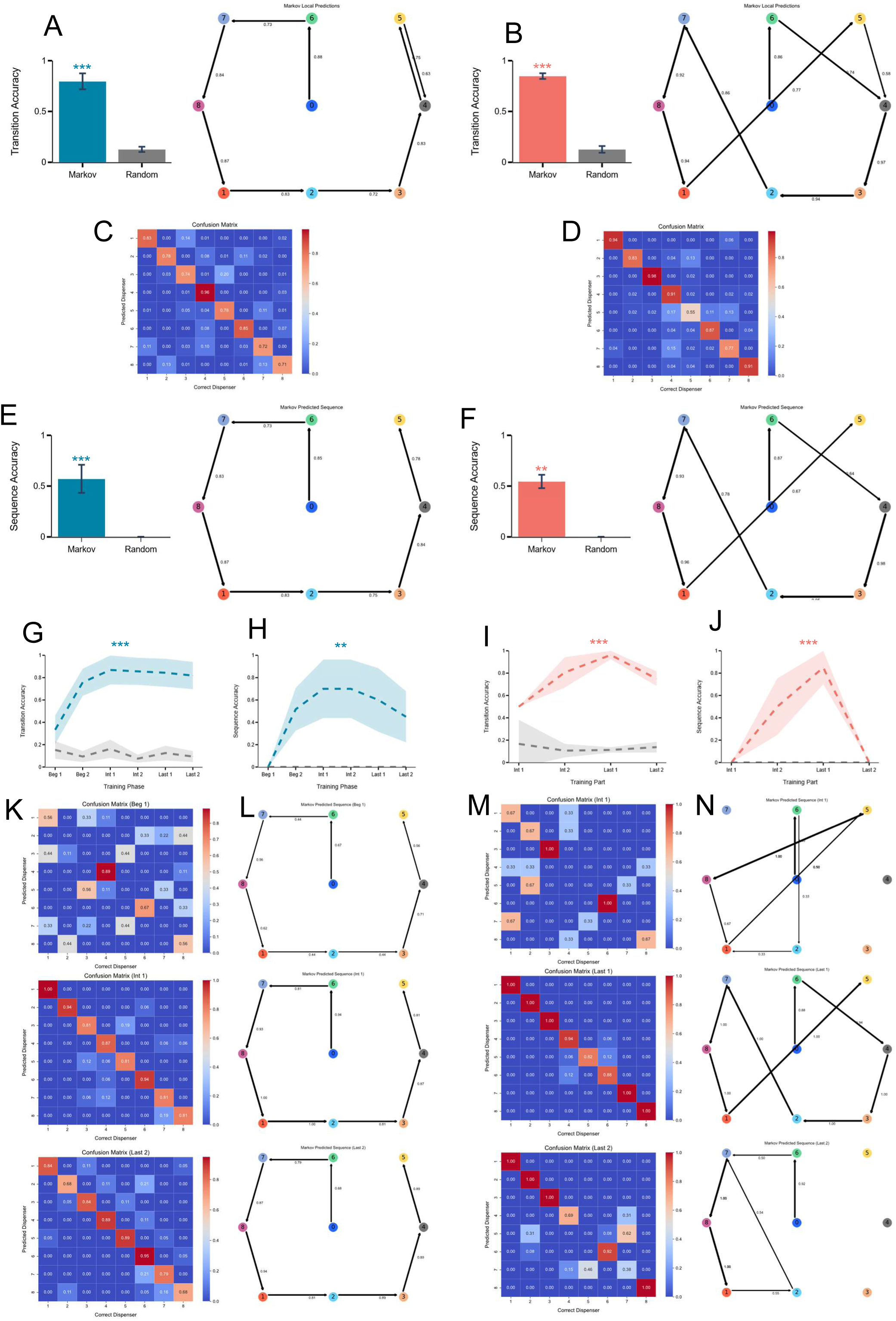
Markov model accuracy at local and global levels across training phases. **A-B. Local transition accuracy and prediction map**. *Left:* Bar plots showing the mean local transition accuracy (across cross-validation folds) for the Markov model and the random baseline. Error bars indicate the 95% confidence interval (CI). Statistical comparisons were made using Wilcoxon signed-rank tests with Bonferroni correction; significant p-values are indicated: ***p < 0.001, **p < 0.01, *p < 0.05. *Right:* Directed graphs depicting the most likely next transition from each dispenser based on transition probabilities estimated from training data. Nodes represent dispensers positioned according to their physical layout; arrow thickness reflects transition probability. Animal E (panel A), Animal F (panel B). **C-D. Normalized confusion matrices for local predictions.** Confusion matrices showing predicted versus actual dispenser choices. Rows are normalized to reflect prediction distributions for each true dispenser. Diagonal values indicate correct predictions. Warmer colors represent higher frequencies. Animal E (panel C), Animal F (panel D). **E-F. Sequence prediction accuracy and transition maps.** *Left:* Bar plots showing mean sequence accuracy across full trials for the Markov model and the random baseline. Error bars represent 95% CIs. Wilcoxon signed-rank tests with Bonferroni correction were used for significance testing. *Right:* Directed graphs illustrating the most probable sequences of up to eight transitions predicted by the Markov model, starting from state 0. Nodes correspond to dispensers, edges indicate predicted transitions. Animal E (panel E), Animal F (panel F). **G-I. Local transition accuracy across training phases.** Line plots showing phase-wise Markov model accuracy (dashed lines) compared to the random baseline. Points correspond to mean accuracy per phase; shaded areas indicate 95% CIs. Kruskal–Wallis tests with Dunn’s post-hoc comparisons assessed accuracy changes over phases; Wilcoxon signed-rank tests compared models within phases (all Bonferroni-corrected). Animal E (panel G), Animal F (panel I). **H-J. Sequence prediction accuracy across training phases.** Same conventions as in panels G and I. Markov model sequence prediction accuracy is shown over training phases alongside the random baseline. Animal E (panel H), Animal F (panel J). **K-M. Confusion matrices across training phases.** Confusion matrices for local predictions at three training stages: early (Beg 1: sessions 1–5), intermediate (Int 2: sessions 16–20), and late (Last 2: sessions 26–30). Color scale indicates prediction frequency per actual dispenser. Animal E (panel K), Animal F (panel M). **L-N. Sequence prediction maps across training phases.** Directed graphs showing the most probable full sequences predicted by the Markov model across three training phases (Beg 1, Int 2, Last 2). Nodes represent dispensers; arrows indicate predicted transitions. Animal E (panel L), Animal F (panel N).

Together, these findings indicate that while individual transitions during successful trials are highly predictable, full sequences retain flexibility. The Markov model accurately predicted local transitions but showed reduced accuracy for complete sequences, suggesting that while dominant transition patterns existed, sequence execution was variable across error-free trials.

### Predictability of local and global transition patterns evolves dynamically across training phases

While the Markov model revealed both local predictability and global flexibility in error-free trials, how these properties evolved over training remained to be determined. We examined the evolution of transition predictability across training phases by evaluating the Markov model’s performance in predicting local transitions and complete sequences within error-free trials. The model’s accuracy was compared to a random baseline model in each phase. Kruskal-Wallis tests assessed phase-wise changes, while Wilcoxon signed-rank tests compared model accuracy against baseline within phases (Bonferroni-corrected).

Error-free trials first emerged at different phases in each animal (Animal E: Beg 1, Animal F: Int 1), but once present, both animals showed a similar trajectory in terms of local transition predictability (**Fig. 5G,K, I,M**).

In early phases, the model did not outperform the random baseline (Animal E: 33.3% ± 12.9%, p > 0.05, Animal F: 50.0% ± 0.0%, p > 0.05), indicating initially unstructured transitions. Over time, accuracy improved significantly (Kruskal-Wallis: Animal E: H = 22.3, p < 0.001, Animal F: H = 16.7, p < 0.001), consistently exceeding the random baseline (p < 0.05 for all comparisons), and peaking at 86.8% ± 12.9% (Animal E) and 96.3% ± 3.7% (Animal F). Accuracy then declined slightly in later phases.

We then evaluated global sequence predictability (**Fig. 5H,L, J,N**). Initially, the model failed to predict any full sequences (Animal E: 0%, Animal F: 0%), reflecting highly variable behavior. Sequence accuracy increased significantly with training (Kruskal-Wallis: Animal E: H = 17.8, p < 0.01, Animal F: H = 22.5, p < 0.001), peaking at 70% ± 26.1% (Animal E, vs random baseline, p < 0.01) and 85% ± 14.96% (Animal F, vs random baseline, p<0.01), before sharply declining in the final phase (Animal E: 45% ± 22.8%, Animal F: 0%).

These findings reveal a dissociation in the dynamics of navigation structure: while local transitions became increasingly stereotyped over time, global sequence patterns fluctuated, reflecting the persistence of navigation flexibility despite local consolidation.

### Sequences of dispenser visits are variable but share core transitions

To further understand the global-level variability observed in successful trials, we examined the structure and recurrence of complete sequences (**Fig. 6A**). Animal E used 22 unique sequences, and Animal F 11, but these were not used equally: a dominant sequence accounted for 65.1% of successful trials in Animal E and 56.5% in Animal F. To quantify this variability, we computed the normalized Shannon diversity index (range: 0-1), which captures both the number and distribution of unique sequences. The resulting values (Animal E: 0.59, Animal F: 0.66) confirmed moderate variability, indicating that while animals relied heavily on a dominant path, they also explored alternative solutions.

**Figure 6:**
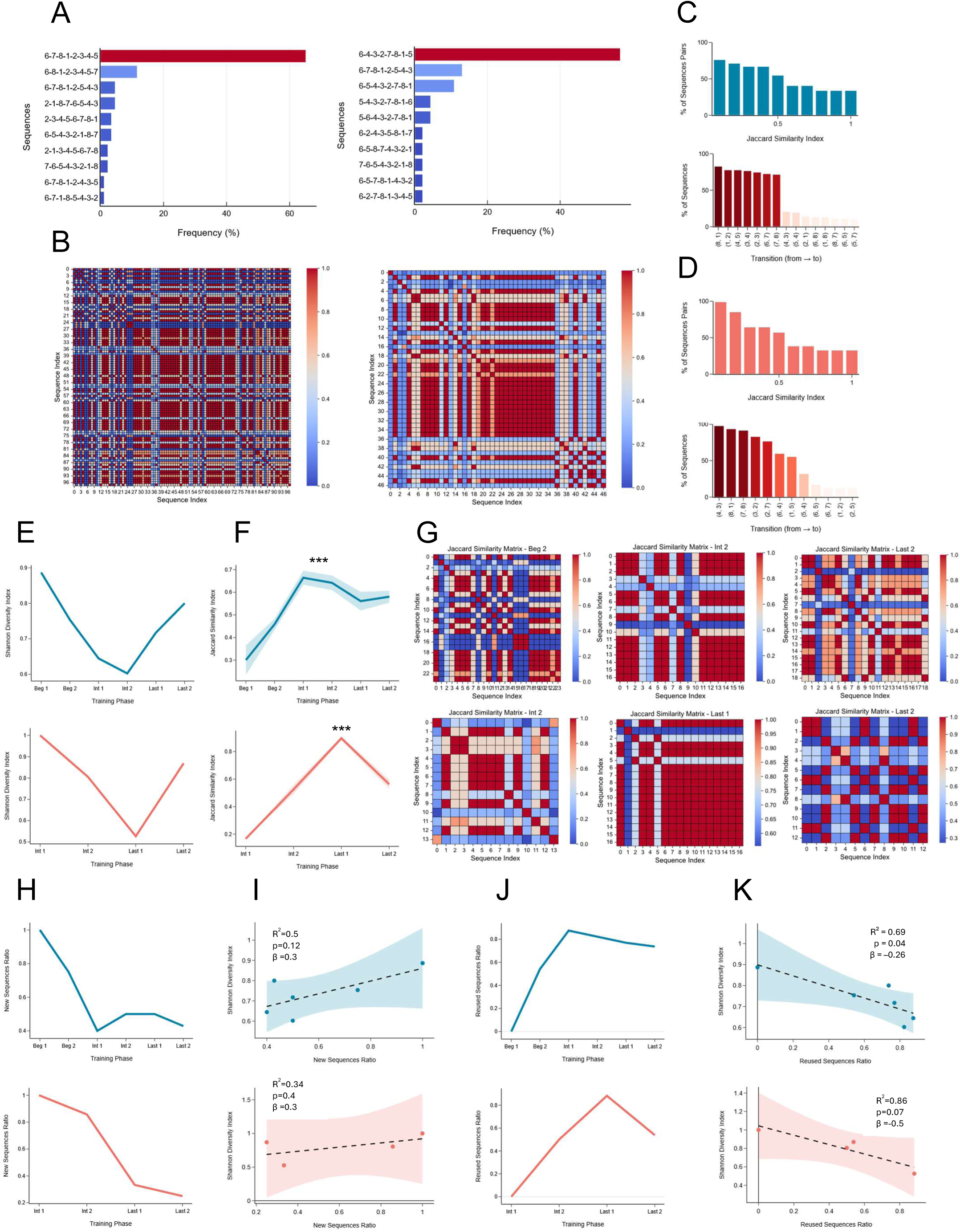
Analysis of sequence diversity, similarity, and strategy reuse across training phases. **A. Top 10 unique sequences in error-free trials.** Horizontal bar plots showing the most frequently used transition sequences during error-free trials. Sequences are sorted by decreasing frequency, with percentages indicating their relative occurrence. *Left*: Animal E; *Right*: Animal F. **B. Sequence similarity (Jaccard index).** Heatmaps displaying pairwise Jaccard similarity between all error-free sequences. Warmer colors indicate greater structural similarity (1 = identical, 0 = no overlap). *Left*: Animal E; *Right*: Animal F. **C.** *Top*: **Pairwise sequence similarity distribution.** Bar plot showing the percentage of sequence pairs with Jaccard similarity equal to or greater than a range of thresholds. This illustrates the degree of structural redundancy or diversity across error-free sequences for Animal E. *Bottom*: **Most frequent transitions.** Bar plot showing the frequency of individual transitions (dispenser-to-dispenser moves) across all error-free sequences. Color intensity reflects recurrence, with darker red bars indicating higher frequency. **D.** Same analyses as in panel C, shown for Animal F. **E. Evolution of sequence diversity**. Line plots showing normalized Shannon diversity index across training phases, with higher values indicating greater sequence variability. *Top*: Animal E; *Bottom*: Animal F. **F. Sequence similarity across training phases.** Line plots showing the evolution of mean Jaccard similarity (± SEM) across training phases. Asterisks indicate significant differences (Kruskal–Wallis test with Bonferroni-corrected Dunn’s post hoc), *** p < 0.001. *Top:* Animal E; *Bottom*: Animal F. **G. Phase-specific similarity matrices.** Heatmaps showing pairwise Jaccard similarity between all error-free sequences within individual training phases. These plots highlight structural convergence in behavior over time. *Top*: Animal E; *Bottom*: Animal F. **H. Exploration of new strategies across training phases.** Line plots showing the evolution of the New Sequence Ratio (NSR), defined as the proportion of sequences in each phase that were not observed in any preceding phase. Higher values indicate greater exploration. *Top*: Animal E; *Bottom*: Animal F. **I. Association between sequence novelty and diversity.** Scatter plots showing linear regressions between NSR and the normalized Shannon diversity index across training phases. Dashed lines represent regression fits; shaded areas indicate 95% confidence intervals. R^2^, p-values, and regression slope (β) quantify the strength and direction of the association. *Top*: Animal E; *Bottom*: Animal F. **J. Reuse of past strategies across training phases.** Line plots showing the evolution of the Reused Sequence Ratio (RSR), defined as the proportion of sequences in each phase repeated from earlier phases. Higher values indicate greater reliance on previously learned sequences. *Top*: Animal E; *Bottom*: Animal F. **K. Association between sequence reuse and diversity.** Scatter plots showing linear regressions between RSR and the normalized Shannon diversity index across training phases. Dashed lines represent regression fits; shaded areas indicate 95% confidence intervals. R^2^, p-values, and regression slope (β) quantify the strength and direction of the association. *Top*: Animal E; *Bottom*: Animal F.

To probe structural similarity across sequences, we extracted all transitions (pairs of consecutive dispensers) and computed the Jaccard similarity index (range: 0-1), between sequences(**Fig. 6B-D**). For Animal E, the mean Jaccard similarity was 0.54 ± 0.006. A majority of sequence pairs (54.7%) shared at least 50% of their transitions, 40.4% fell between 60-70% similarity, and 33.8% were identical. Notably, 7 transitions appeared in over 70% of sequences, reflecting recurring motifs. For Animal F, the pattern was comparable: mean Jaccard similarity reached 0.58 ± 0.01, with 56.9% of sequence pairs sharing ≥50% of transitions, 38.4% between 60-70%, and 32.6% identical. Five transitions were present in over 75% of sequences.

These findings indicate that, while animals occasionally used alternative routes to complete the task, most sequences converged around a core set of transitions. This suggests that even when behavior appears variable at the sequence level, it is underpinned by consistent structural motifs.

### Dynamics of sequence diversity and similarity

We next asked when this structural consistency emerged during training. Specifically, we examined how sequence diversity and similarity evolved across training phases to identify when animals began to converge on consistent behavioral motifs.

We first analyzed changes in sequence diversity using the normalized Shannon diversity index (**Fig. 6E**). Both animals showed a non-linear U-shaped trajectory across training. To quantify this pattern, we fitted a quadratic model to the diversity values (**Fig. S5A**). For Animal E, the model accounted for a large proportion of the variance (R^2^ = 0.96, p < 0.01), confirming the significance of the U-shape. For Animal F, the fit was also strong (R^2^ = 0.86) and outperformed a linear model (R^2^ = 0.4), although it did not reach significance due to the limited number of phases with error-free trials (p = 0.3). Nonetheless, the improvement in fit suggests that a similar non-linear trend likely applies to both animals.

To further probe this trajectory, we examined how the use of the dominant sequence changed across training phases (**Fig. S5B**). In both animals, its frequency followed an inverted U-shaped pattern, initially low, rising to a peak mid-training, then declining in later phases. This trend paralleled changes in diversity: dominant sequence frequency was negatively correlated with the Shannon index (**Fig. S5C**, Animal E: r = -0.95, p < 0.01, Animal F: r = -0.95, p = 0.05), confirming that increased reliance on a single sequence coincided with reduced sequence diversity.

We then assessed how sequence similarity evolved by computing the Jaccard index between all sequence pairs within each training phase (**Fig. 6F-G**). In both animals, similarity increased significantly over training (Kruskal-Wallis, Animal E: H = 50.38, p < 0.001, Animal F: H = 102.74, p < 0.001), peaking in Int 1 (Animal E: 0.66 ± 0.03) and Last 1 (Animal F: 0.89 ± 0.013). Although similarity declined in the final phase (Dunn tests, Animal E: Int 1 vs Last 2, p < 0.05, Animal F: Last 1 vs Last 2, p < 0.001), it remained significantly higher than in early training (Animal E: Beg 1 vs Last 2, p < 0.001, Animal F: Int 1 vs Last 2, p < 0.05). These results indicate that, as training progressed, the sequences used in error-free trials within each phase increasingly shared common transitions, reflecting a growing consistency in the animals’ transition patterns.

To assess whether this structural consistency persisted across phases, we computed the similarity between sequences from different training periods. In both animals, late-phase sequences shared a substantial proportion of transitions with earlier ones (**Fig. S5D),** indicating that core transitions were not confined to a specific phase but remained stable over time.

Together, these results show that structural consistency in behavioral sequences emerged progressively over the course of training. Early phases were characterized by high variability and low similarity between sequences, suggesting that animals initially deployed a broad set of behavioral solutions. In later phases, they increasingly converged on dominant, structurally similar trajectories, reflecting a gradual stabilization of their navigation patterns. Although sequence diversity rose again in the final phase, core transition motifs remained stable both within and across phases. This temporal persistence suggests that animals increasingly relied on a stable set of transitions, rather than transient, phase-specific strategies.

### Marmosets’ shift from novel to reused sequences across training phases

The observed changes in sequence diversity and similarity indicate that structural motifs became increasingly consistent within and across training phases. However, these patterns do not reveal how such stability emerged. Was this convergence driven by the continuous generation of new but structurally similar sequences, or by the repeated use of previously established ones? To address this, we sought to characterize how animals balanced exploration and exploitation over training, that is, to what extent variability arose from the search for novel strategies versus the selective reuse of learned ones.

We quantified the relative contributions of novel and repeated sequences over time using two complementary metrics. The New Sequence Ratio (NSR) captured the proportion of unique sequences in each phase that had not been used previously, while the Reused Sequence Ratio (RSR) quantified how often sequences from earlier phases reappeared. High NSR values reflect active exploration of novel solutions, whereas high RSR values indicate growing reliance on familiar trajectories retrieved from past experiences.

Both animals displayed a similar progression (**Fig. 6H,J**). Early training phases were marked by high NSR and low RSR values, consistent with broad exploration. Over time, NSR progressively declined while RSR increased, indicating a gradual shift toward the reuse of previously successful sequences and reflecting a growing tendency to repeat previous sequences within each training phase.

To determine which behavioral metric best explained within-phase sequence diversity, we fitted separate linear models using each metric as a predictor of Shannon diversity index (**Fig. 6I,K**). The RSR emerged as the strongest predictor of variability, particularly in Animal E (R^2^ = 0.69, p = 0.04, β= -0.26). In Animal F, a similar negative trend was observed (R^2^ = 0.86, p = 0.07, β = -0.51), though it did not reach statistical significance, likely due to the small sample size (n = 4). In contrast, the NSR showed a weaker association with diversity (Animal E: R^2^ = 0.5, p = 0.12, β =0.3, Animal F: R^2^ = 0.34, p = 0.4, β =0.3), suggesting that exploration of novel solutions contributed less to variability.

Together, these findings reveal that, over the course of training, animals progressively shifted from generating new solutions to flexibly reusing learned ones, reflecting a dynamic balance between exploration and exploitation. Moreover, behavioral diversity appeared more strongly shaped by reuse of previous sequences than by the continued generation of novel ones.

## Discussion

In this study, we examined how freely moving marmosets progressively structured their navigation through repeated experience in a semi-naturalistic task. Without instruction or shaping, animals transitioned from exploratory to organized behavior, progressing through discrete states that reflected engagement and execution quality. Learning occurred selectively within engaged states, marked by reduced errors and the emergence of efficient navigation patterns. Over time, animals consolidated a stable repertoire of locally stereotyped transitions, while preserving global flexibility through the recombination of familiar elements. These findings offer a rare behavioral framework for understanding how structured yet flexible navigation strategies emerge through self-guided experience in non-human primates (NHPs).

### A naturalistic paradigm to study spatial memory in marmosets

Unlike classical marmoset tasks that investigate memory using touchscreens, constrained locomotion, or simplified mazes ^19–22^, our paradigm involves naturalistic, three-dimensional navigation, enabling spontaneous and ecologically relevant behavior. This setting offers a powerful opportunity to study how spatial memory systems operate under real-world-like conditions. The value of naturalistic paradigms is increasingly recognized in neuroscience, as core functions like memory, learning, and decision-making are more richly expressed and adaptively deployed in unconstrained, ecologically valid contexts ^2,23–25^. Our task embraces this shift by combining controlled contingencies with the freedom to observe self-guided learning. Without instructions or shaping, animals progressively organized their navigation through repeated experience, developing efficient and flexible strategies. This unsupervised structuring of behavior reveals how cognitive patterns emerge from free interaction with a complex environment. Interestingly, the navigation patterns observed in our task resemble spatial behaviors reported in wild marmosets^17,18^, further supporting the ecological validity of our framework for studying naturalistic spatial navigation.

### Spatial learning emerges from distinct states of engagement

A key challenge in minimally constrained settings is to distinguish genuine learning from fluctuations in motivation or task engagement. In our dataset, animals exhibited strong trial-by-trial variability, which we leveraged to identify discrete behavioral states using unsupervised clustering. Two of these states reflected active task engagement, differing in execution quality, while the others captured disengagement. Crucially, only the engaged states changed over time: efficient execution increased while error-prone behavior declined, indicating that learning occurred selectively within periods of engagement. Disengaged states, by contrast, remained stable across sessions. This finding underscores the importance of dissociating learning-related changes from spontaneous variability in unconstrained settings. Rather than averaging across heterogeneous trials, isolating periods of genuine engagement allows for a more accurate characterization of memory-guided learning. This approach aligns with recent advances in computational neuroethology, which use unsupervised models to uncover latent structure in natural behavior^26,27^.

### Refinement of spatial working and reference memory under self-guided learning

During active engagement, we observed the refinement of different memory systems, in the absence of instruction or corrective feedback. The decline in repetition and rule errors over time suggests improved spatial working memory and better internalization of task structure, consistent with previous evidence that marmosets possess robust working memory capacities^20^. In parallel, the increasing reuse of previously successful sequences, quantified by the Reused Sequence Ratio (RSR), pointed to the gradual consolidation of spatial reference memory, consistent with recent findings showing that marmosets can form stable long-term spatial memories^28^. Together, these patterns suggest that animals combined short-term tracking and long-term spatial knowledge to guide navigation, highlighting the co-development of complementary memory systems in our semi-naturalistic setting.

### Trajectory structuring reveals the behavioral signature of memory consolidation

How this refinement of memory shaped behavior became evident in the structure of movement trajectories. With experience, animals shifted from dispersed exploration to more selective and reproducible transitions between dispenser locations. This was captured by a progressive decrease in both global and local entropy, increased similarity across transition matrices, and reduced Euclidean distances over sessions. Sequence similarity analyses confirmed a convergence toward shared paths, while a first-order Markov model revealed increasing predictability of local transitions, consistent with the stabilization of specific movement motifs. This structuring emerged within a shared transition repertoire between engaged states: both efficient (Cluster 0) and error-prone (Cluster 1) trials relied on similar local patterns but differed in their consistency. The shift from Cluster 1 to Cluster 0 across training suggests that learning involved not only discovering efficient routes but increasingly stabilizing their use. These results indicate that spatial memory consolidation was behaviorally expressed in the form of organized, repeatable movement patterns. While similar dynamics have been reported in rodents^29–31^, our study provides the first longitudinal evidence that such trajectory-level consolidation can emerge through self-guided learning in freely moving NHPs.

### Structured yet flexible navigation suggests route-based strategies anchored in topological representations

The emergence of structured navigation routines raises a deeper question: what kind of spatial strategies support this organization? Classically, two broad strategies have been proposed: route-based navigation, relying on learned movement sequences or landmark associations, and map-like representations, supporting flexible planning based on internal models of space. Distinguishing between these strategies sheds light not only on behavioral algorithms but also on the underlying neural mechanisms^9–12^. To probe the nature of the marmosets’ strategy, we used a first-order Markov model to assess how well transitions between dispensers could be predicted from the immediately preceding location. The model’s high and increasing accuracy over training suggests a growing reliance on stereotyped local transitions, consistent with the use of a route-based strategy. These dominant transitions appear to have been reinforced through repeated experience and remained stable across sessions, indicating the consolidation of a temporally persistent movement repertoire. However, animals did not rigidly replay full routes. When the model was applied to entire sequences, its performance dropped substantially. Sequence similarity analyses revealed that animals generated variable trajectories composed of overlapping transitions, rather than repeating fixed paths. Spatial entropy analyses further showed that, although global structure increased with training, some locations retained high local variability, suggesting flexible decision points embedded within otherwise stable routines. This coexistence of reliability and flexibility is a hallmark of topological representations, in which navigation relies on a graph-like network of spatial relationships rather than on fixed sequences^11,12^. Altogether, our findings point toward a hybrid strategy: through repeated experience, marmosets assembled a structured set of familiar route segments, while preserving the flexibility to recombine them in varying orders. This organization mirrors patterns observed in wild marmosets^17,18^ and aligns with theoretical models of topological navigation in rodents and humans^10,12,32^, where local regularities support efficient navigation while preserving global adaptability.

### Dynamic shifts in sequence use reflect an exploration-exploitation trade-off

A central feature of our findings is the evolving balance between behavioral stability and variability over training. Early in learning, animals explored broadly, generating diverse and unstructured sequences. With experience, they converged on a core set of solutions, marked by increased reuse of familiar transitions. Yet this consolidation did not lead to rigidity: in later stages, animals flexibly recombined known elements into novel configurations, maintaining variability within a stabilized scaffold. This was reflected in a U-shaped trajectory of sequence diversity and Markov model predictability, a pattern consistent with dynamic exploration-exploitation trade-offs as described in decision-making and learning domains^33–35^. Such dynamics typically involve broad early exploration, followed by exploitation of reinforced strategies, and a late reintroduction of flexibility to support adaptation or fine-tuning. Importantly, this renewed variability was not driven by the discovery of novel paths: the New Sequence Ratio (NSR) declined steadily, while the Reused Sequence Ratio (RSR) increased and strongly predicted sequence diversity. This suggests that late-stage variability reflected the flexible deployment of previously consolidated patterns, rather than novelty-seeking. These findings extend reinforcement learning frameworks in NHPs beyond isolated choices^36–38^, revealing how structured spatial behavior can emerge, stabilize, and later reorganize through self-guided learning. They also resonate with recent theories emphasizing internal planning and information sampling^39^, suggesting that primates may balance efficiency and flexibility by proactively recombining known solutions.

### Neural correlates of memory-guided navigation in primates

The behavioral transformations observed in our task, ranging from memory refinement to the consolidation of structured yet flexible trajectories, likely reflect the joint contribution of multiple neural systems involved in memory, decision-making, and strategy selection. Classically, spatial learning has been described as a dual-system process: the hippocampus supports flexible, relational representations, while the striatum underlies reinforcement-based route acquisition and automatization ^9,40^. However, growing evidence in rodents and primates supports a more integrated view, in which these systems interact dynamically rather than operating in strict competition^41–45^. This aligns with our findings: marmosets did not exhibit a linear transition from cognitive to habitual strategies, but rather a dynamic balance between stability and flexibility, consistent with ongoing hippocampal-striatal coordination. This coordination is likely further modulated by the prefrontal cortex, which plays a critical role in integrating contextual goals, maintaining task-relevant information, and regulating the exploration-exploitation trade-off ^46–49^. In our task, the ability to flexibly adapt familiar routes, follow task-relevant rules, and alternate between variability and consolidation suggests the involvement of prefrontal mechanisms in orchestrating behavioral flexibility across timescales, a pattern consistent with integrative models of prefrontal-hippocampo-striatal interaction during spatial navigation ^50^.

While the hippocampo-striatal-prefrontal architecture appears broadly conserved across mammals, key sensorimotor differences shape its implementation in primates. Unlike rodents, which rely heavily on olfaction and path integration, primates, including marmosets, engage in visually guided navigation, combining landmark use with active gaze exploration ^3,51^. These differences are reflected in distinct neural coding schemes: in rodents, hippocampal place cells encode 2D allocentric space ^52^, whereas in primates, hippocampal neurons show strong modulation by gaze direction and viewpoint, supporting more relational, view-dependent spatial representations^53–56^. Notably, such coding has now been demonstrated in freely moving marmosets: hippocampal cells encode 3D position as well as head- and gaze-centered views^57^, linking their visually guided exploration of 3D space to distinct hippocampal dynamics under naturalistic conditions. Together, these findings reinforce the marmoset’s value as a model for studying primate-specific navigation mechanisms. Recent comparative work shows that marmosets share core perceptual visual strategies with macaques and humans^1^. This continuity suggests that the spatial navigation processes observed in marmosets may reflect fundamental features of primate cognition that are less accessible in rodent models.

### Limitations and future directions

While our paradigm enabled a rare longitudinal view of spatial learning in freely moving primates, some limitations should be acknowledged. First, the study was conducted on a small number of animals. However, this constraint is common in primate neuroscience, where methodological and ethical considerations often limit sample size^58,59^. It is, moreover, partly mitigated by the robustness of behavioral patterns across individuals and the richness of within-subject data collected over time. Importantly, small-N longitudinal designs have previously yielded valuable insights into primate cognition, particularly when intra-individual dynamics are the primary focus. Second, our semi-naturalistic setting maximized ecological validity within a laboratory context, but reduced experimental control over the sensory cues guiding navigation. In this open setup, animals could rely on various combinations of visual, proprioceptive, or olfactory information, making it difficult to disentangle their respective contributions. Future studies could address this by selectively manipulating sensory inputs (e.g., masking olfactory cues, modifying visual landmarks) or introducing spatial perturbations to assess cue integration and weighting. Still, this openness allowed animals to develop ethologically relevant strategies that might be obscured in more constrained paradigms, illustrating the trade-off between ecological richness and experimental control. Finally, our study did not include neural recordings, and inferences at the circuit level therefore remain hypothetical. While the behavioral dynamics we observed align with models of hippocampal, striatal, and prefrontal contributions to navigation, direct evidence is needed to confirm how these circuits interact during self-guided learning. Integrating chronic electrophysiological recordings or imaging into this behavioral framework will be essential to uncover how conserved and primate-specific neural dynamics support flexible spatial behavior.

## Conclusion

Our study shows that structured, memory-guided behavior can emerge spontaneously in primates through repeated, self-guided interaction with a complex environment. By combining ecological richness with longitudinal precision, our paradigm reveals how distinct memory systems co-develop to support flexible yet stable navigation, learning processes rarely captured in non-human primates. Rather than forming rigid routes or relying solely on short-term memory, marmosets gradually assembled and flexibly reused a repertoire of spatial motifs embedded in a topological structure. These findings offer a framework for linking naturalistic behavior to neural computations, and open new directions for exploring how sensorimotor constraints shape the neural basis of spatial cognition across species. This work bridges naturalistic ethology and systems neuroscience, and lays the groundwork for integrating circuit-level recordings with the study of structured, adaptive behavior in freely moving primates.

## Supplemental information

Document S1. Figures S1-S5, Tables S1 and S2.

## Acknowledgments

We thank the animal care staff at the Ernst Strüngmann Institute for their assistance in ensuring the welfare of the marmosets. A specific thanks to Colleen Illing for supporting the marmosets’ training. This work was supported by Bundesministerium für Bildung und Forschung (ERANET NEURON 01EW2110).

## Author contributions

Conceptualization: N.E.M., F.L., J.L., Methodology: N.E.M., F.L., Investigation: N.E.M., F.L., Formal Analysis: N.E.M., Technical Support: F.Z.N., D.S., Writing - Original Draft: N.E.M., F.L., Writing - Review & Editing: N.E.M., F.L., F.Z.N., D.S., J.L., Funding Acquisition: J.L., Supervision: J.L. All authors read and approved the final manuscript.

## STAR★Methods

### Key Resources table

| REAGENT or RESOURCE | SOURCES | IDENTIFIERS |
| --- | --- | --- |
| **Deposited data** |  |  |
| Raw behavioral data for all analyses and figures | This study | https://github.com/nadaelma/Marmoset_spatial_navigation |
| **Software and algorithms** |  |  |
| scikit-posthocs | Terpilowski, 2019 | https://github.com/maximtrp/scikit-posthocs |
| statsmodels | Seabold & Perktold, 2010 | https://www.statsmodels.org |
| BORIS (Behavioral Observation Research Interactive Software) | Friard & Gamba, 2016 | https://www.boris.unito.it |
| Custom Python analysis script (including Markov model) | This study | https://github.com/nadaelma/Marmoset_spatial_navigation |
| **Experimental Models: Organisms/Strains** |  |  |
| *Callithrix jacchus* | German Primate Center, Göttingen, Germany | N/A |

### Resource availability

#### Lead contact

Requests for further information and resources should be directed to and will be fulfilled by the lead contact, Nada El Mahmoudi.

#### Materials availability

This study did not generate any new unique reagents.

#### Data and code availability

All data and code to reproduce the main results will be made publicly available on a GitHub repository (https://github.com/nadaelma/Marmoset_spatial_navigation) upon acceptance of the manuscript. We are happy to provide the code upon request during peer review.

### Experimental model and subject details

#### Ethical Statement

All animal procedures in this study were approved by the relevant regional government authority (F149/1012) and adhered to all applicable German and European regulations on animal husbandry and welfare. A small-sample approach was adopted, in line with previous research on spatial behavior in freely moving non-human primates ^20,51,58,60^. This approach allowed for a substantial amount of data to be collected per animal, minimizing the need for a larger sample while maintaining methodological rigor ^59^.

#### Animals and Housing

Two adult male common marmosets (Callithrix jacchus, mean age: 3 years) participated in this study. The animals were pair-housed in a cage measuring 180 cm (H) × 90 cm (W) × 60 cm (D) under a 12-hour light-dark cycle (07:00-19:00). Room temperature was maintained at 24-25°C with 60% relative humidity. The marmosets were obtained from the German Primate Center (Göttingen, Germany). Experimental sessions took place in the morning. In addition to the gum arabic collected during the task, animals were fed a daily meal consisting of pellets, a dietary supplement primarily composed of quark, yogurt, banana and vitamins, and fresh vegetables. Water was provided ad libitum.

### Method details

#### Experimental Apparatus and Data Collection

The experimental setup consisted of a wire mesh enclosure measuring 1.8 m × 1.4 m × 1.6 m (total volume: 4 cubic meters). The wire mesh walls facilitated climbing and allowed the marmosets to navigate freely within the space. The entrance was located in a designated corner, where a tunnel system enabled transition from the transport cage to the setup. To encourage locomotion and exploratory behavior, the enclosure was enriched with wooden sticks, semi-rigid ropes, and suspended enrichment items (**Fig.1A**). Additional environmental cues were introduced around the enclosure, including four white walls with landscape posters, four colored curtains, and large plastic plants.

During each experimental session, rewards were dispensed using custom-made, 3D-printed, remotely controlled dispensers (**Fig. 1A, E**). Each dispenser was calibrated to deliver 3 ml of liquid gum arabic per session, in increments of approximately 0.15 ml per activation. To ensure accurate dispensing, the system was tested and calibrated prior to each session.

Two experimenters conducted the sessions individually, alternating roles between sessions. During each session, the experimenter was positioned at a fixed location at a predetermined distance from the setup, manually scoring dispenser visits and task latencies in real time using BORIS, a well-established behavioral event logging software ^61^. A visit to a dispenser was defined as the marmoset pausing in front of the dispenser and attempting to retrieve the reward. This criterion ensured that only purposeful interactions with the dispenser were recorded as visits. Additionally, a wide-angle lens camera was mounted above the setup, continuously recording the sessions for further analysis.

#### Behavioral Training

##### Habituation

Before the experiment, the marmosets underwent a habituation phase to ensure they did not exhibit anxiety and displayed natural exploratory behavior in the setup. This process took place over multiple sessions and was conducted by both experimenters, who had prior experience training the animals. As a result, the marmosets were well-acquainted with both experimenters before the start of behavioral testing. Initially, the two marmosets were trained to be comfortably transported from the animal facility to the laboratory in a transport cage. They were then introduced together into the setup, where they could explore and discover hidden rewards (e.g., marshmallows) placed in various locations. In the final habituation phase, only one marmoset entered the setup at a time, while the second remained in the transport cage nearby, maintaining visual contact. Initially, the separation lasted only a few minutes but gradually increased until the marmoset could remain alone for at least 30 minutes without signs of discomfort. Throughout this process, the experimenter monitored welfare by assessing vocalizations and behaviors based on established ethological criteria for captive marmosets ^16^.

##### Home Dispenser Training

Since the marmosets explored the setup freely, they needed to be trained to voluntarily move to the center of the apparatus when required, ensuring that each trial started from a standardized position during the spatial working memory task. To achieve this, a food dispenser (i.e the “Home Dispenser (HD)”) was installed upside down at the center of the ceiling (**Fig.1E**). The HD contained liquid gum arabic and was equipped with a blue LED and a buzzer, providing visual and auditory cues upon activation. Using positive reinforcement, the monkeys were trained to associate these cues with the HD activation and to reach it promptly.

Each marmoset was tested individually. At the start of each trial, the HD was activated for ∼3 seconds, signaling the monkey to move to the center. The subject had up to 60 seconds to reach the HD for the trial to be considered successful. If the monkey failed to do so within this timeframe, the trial was marked as unsuccessful. A session was terminated if the monkey failed to complete two consecutive trials. Performance was evaluated based on the number of successful trials per session and the latency to reach the HD. Training was considered complete once the monkey achieved over 80% successful trials in two consecutive sessions. On average, each session consisted of 20.4 ± 1.19 trials for Animal E and 16.05 ± 1.2 trials for Animal F. To reach the training completion criterion, Animal E required 10 sessions, while Animal F needed 18 sessions.

#### Spatial memory task (Behavioral testing)

The experiment assessed spatial working memory by requiring marmosets to retrieve rewards from eight dispensers while remembering which ones they had already visited.

During the testing phase, eight additional dispensers were introduced, placed in fixed locations along the perimeter of the setup to ensure uniform spatial coverage (**Fig.1A**). Specifically, three dispensers were placed equidistantly along each of the two long sides, while one dispenser was positioned at the midpoint of each short side. This configuration ensured consistent spatial cues and a uniform distribution of reward locations across trials.

At the start of each trial, the Home Dispenser (HD) was activated, prompting the monkey to move to the center of the apparatus within 60 seconds. Once the monkey reached the HD, all eight dispensers were remotely activated simultaneously (**Fig.1B**)., each releasing ∼0.15 ml of gum arabic as a reward. The food was concealed within the spout of each dispenser and not visible to the monkey, requiring them to actively search for and retrieve it. The reward remained available until collected.

To successfully complete the trial, the monkey had to visit all eight dispensers at least once, retrieving the reward from each. The trial ended upon the first visit to the eighth unique dispenser, regardless of whether the monkey revisited some dispensers along the way. However, if the monkey failed to visit all eight dispensers within 10 minutes, the trial was automatically terminated. After trial completion or timeout, the HD was reactivated, signaling the start of the next trial.

Each monkey underwent 30 training sessions, with an average of 7.96±0.4 trials for Animal E and 8±0.4 trials for Animal F per session. A session was terminated if the monkey failed to complete two consecutive trials.

### Data analysis

All data analyses were performed in Python using custom-made scripts. Given the single-subject design, non-parametric tests were preferred. When needed, parametric analyses were conducted with model assumptions systematically verified, and robust standard errors applied when necessary. Bonferroni corrections were used to control for multiple comparisons. Statistical significance was set at α = 0.05, with additional thresholds reported for p < 0.01 and p < 0.001 where relevant.

#### Spatial distribution of visits and omissions

To assess whether the marmosets interacted equally with all dispensers, we analyzed the number of visits and omissions for each of the eight dispensers. We first used Chi-squared tests to compare the observed visit and omission distributions to an expected uniform distribution, testing whether interaction patterns were evenly distributed. If a significant deviation was detected, binomial tests were conducted for each dispenser to identify which ones had visit frequencies that significantly differed from chance. A Bonferroni correction was applied to control for multiple comparisons.

#### Error Analysis and Regression Across Sessions

To assess error patterns across training sessions, errors were categorized into three types: rule errors (returning to the home dispenser during a trial), repetition errors (revisiting a dispenser within the same trial), and omission errors (failing to visit a dispenser). The total number of errors per trial was computed and averaged per session to obtain the mean number of errors per session and the standard error of the mean (SEM).

A linear regression analysis was performed to evaluate changes in the mean number of errors over time, with session number as the independent variable. The slope of the regression line was examined to determine whether errors decreased with training. Model fit was assessed using R-squared values, and confidence intervals were computed to visualize prediction uncertainty. To ensure the validity of the regression model, key assumptions were tested: normality of residuals was assessed using the Omnibus test, independence of errors was verified with the Durbin-Watson statistic, and homoscedasticity was examined using the Breusch-Pagan test. Heteroscedasticity, if detected, was addressed by computing robust standard errors.

To further investigate factors influencing error rates, a multiple linear regression was conducted with session number, number of trials, number of days since the last session, and experimenter identity as independent variables. Model assumptions were systematically tested to ensure the validity of the regression analysis. Normality of residuals was assessed using the Shapiro-Wilk test, homoscedasticity was evaluated with the Levene test, and multicollinearity was examined through the Variance Inflation Factor (VIF). Ordinary Least Squares (OLS) regression with heteroscedasticity-robust standard errors was applied to account for variance inconsistencies. To control for multiple comparisons, p-values were adjusted using Bonferroni corrections.

#### Error Type Dynamics and Correlation Analysis

To examine the evolution of error types across training, we calculated the mean number and SEM of rule errors, repetition errors, and omission errors per session. A Kruskal-Wallis test was used to assess whether these error distributions differed significantly across sessions, followed by Dunn’s post hoc test with Bonferroni correction for multiple comparisons. To explore relationships between error types, two correlation matrices were computed: one at the session level, where values were averaged per session, and one at the trial level, using individual trial data. Both matrices included total errors, total number of visits, rule errors, repetition errors, and omission errors. Pearson’s correlation coefficients were used to assess linear associations, with Bonferroni correction applied to control for multiple comparisons.

#### Clustering analysis

##### Clustering procedure

We conducted a k-means clustering analysis to identify distinct behavioral patterns in marmoset task execution based on omission errors (failing to visit a dispenser) and total visits to dispensers. These two variables were selected to capture complementary aspects of task engagement, with omission errors reflecting disengagement and total visits indicating active task participation. Prior to clustering, the data were standardized to ensure equal weighting of both variables. The optimal number of clusters was determined by balancing silhouette scores and inertia, which assess cluster separation and within-cluster cohesion, respectively. Each trial was then assigned to a cluster based on proximity to the cluster centroids, representing the mean omission errors and total visits per cluster.

##### Statistical Comparison Between Clusters

To assess differences in task performance among the clusters identified by K-Means clustering, we analyzed several performance variables, including trial duration, total number of visits, and error types (rule errors, repetition errors, and omission errors). A Kruskal-Wallis test was conducted to determine whether these variables differed significantly across clusters. If significant differences were detected (p < 0.05), Dunn’s post-hoc test was performed for pairwise comparisons, with a Bonferroni correction applied to control for multiple comparisons.

##### Regression Analysis of Cluster Proportions Across Sessions

A linear regression analysis was performed to evaluate changes in the proportion of trials assigned to each cluster over training, with session number as the independent variable. Separate ordinary least squares (OLS) regression models were fitted for each cluster. The slope of the regression line was examined to determine whether the proportion of trials in each cluster changed significantly over time. Model fit was assessed using R-squared values, and confidence intervals were computed to visualize prediction uncertainty. To ensure the validity of the regression models, normality of residuals was assessed, independence of errors was verified using the Durbin-Watson statistic, and homoscedasticity was examined using the Breusch-Pagan test. When necessary, heteroscedasticity was addressed by computing robust standard errors.

#### Analysis of performance over the first eight choices

To assess task performance, we analyzed the success rate over the first eight choices, defined as the proportion of correct responses within these first eight selections. A response was considered correct if the marmoset selected a dispenser that had not been visited previously in the trial. This analysis was performed in two contexts: (1) comparison between clusters 0 and 1, and (2) evolution of performance across training phases.

##### Cluster Performance Analysis

To determine whether performance within each cluster differed from what would be expected under random selection, we computed the expected random performance assuming choices were made randomly among nine available options. Since the number of remaining correct options decreases at each step, the probability of selecting a correct option follows this sequence:

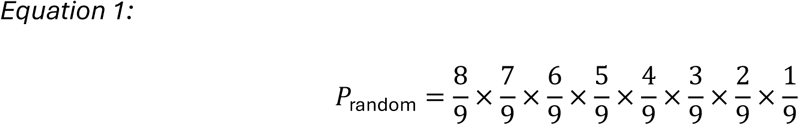

The expected average success rate was then computed as:

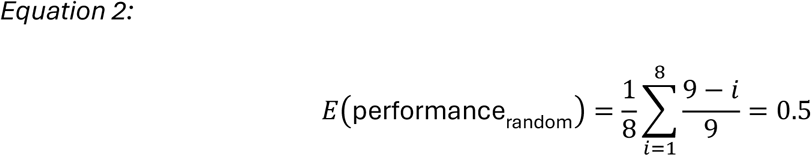

Thus, under random selection, the expected success rate is 50%, meaning that, on average, a marmoset would make 4 correct choices out of 8 by chance. This value served as a baseline for evaluating observed performance in each cluster.

For each cluster, we calculated the mean success rate and SEM. To assess whether cluster performance significantly deviated from the random expectation, we performed a Wilcoxon signed-rank test against the 50% baseline, with Bonferroni correction applied for multiple comparisons. Pairwise comparisons between clusters were conducted using a Mann-Whitney U test.

##### Performance Evolution Across Training Phases

To evaluate changes in performance over training, we divided the training sessions into six phases, each consisting of five sessions: the first ten sessions were divided into Beg 1 (S1-S5) and Beg 2 (S6-S10), the ten intermediate sessions into Int 1 (S11-S15) and Int 2 (S16-S20), and the last ten sessions into Last 1 (S21-S25) and Last 2 (S26-S30). For each phase, we computed the mean performance rate and SEM. Performance rates were compared across these six phases using a Kruskal-Wallis test to determine whether task performance significantly improved over time.

#### Transition Patterns Analysis

##### Cluster Analysis

###### Transition Probability Computation

Each trial was represented as a sequence of visits to dispensers. Transition probabilities between locations were calculated by determining the frequency of transitions from one dispenser to another. The probability of transitioning from dispenser *i* to dispenser *j* was defined as:

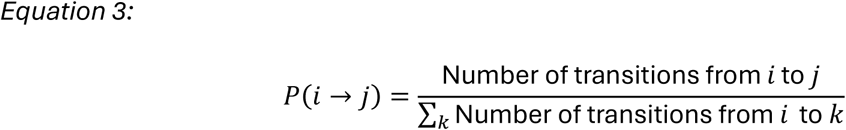

Where *P*(*i* → *j*) is the probability of transitioning from dispenser *i* to dispenser *j*, and the denominator is the total number of transitions starting from dispenser *i* to any possible dispenser (the sum over all possible dispensers *k*).

To ensure comparability across clusters, we considered all possible transitions-including those with zero occurrences-and then normalized the transition probability matrix on a row-by-row basis. In other words, for each dispenser (row), we scaled the counts so that the sum of the probabilities for all transitions starting from that dispenser equals one.

###### Global Entropy and Global Transition Diversity

For the calculation of global entropy and global transition diversity, the transition probability matrix was further normalized globally. That is, after computing the row-wise transition probabilities for each dispenser, we rescaled the entire matrix so that the sum of all transition probabilities within each cluster equals 1. This global normalization allows for a direct comparison of transition patterns between clusters.

We calculated global entropy based on Shannon entropy’s formula:

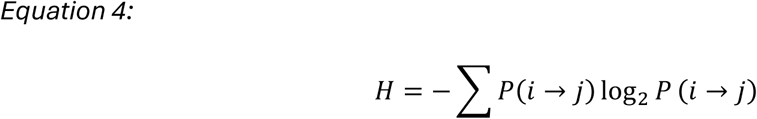

Where *P*(*i* → *j*) represents the probability of transitioning from dispenser *i* to dispenser *j*. In this context, global entropy quantifies the overall unpredictability of transitions across all dispensers: higher entropy values indicate that transitions are more varied and less predictable, while lower values suggest a more structured, stereotyped pattern dominated by a few transitions.

In addition, we calculated global transition diversity using Simpson’s Diversity Index to assess the evenness of transition probabilities between dispensers. The index was computed as:

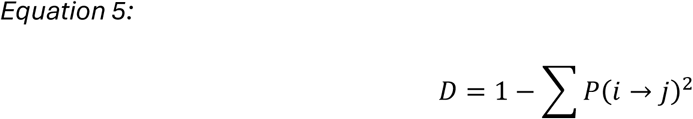

In the context of transition patterns, global transition diversity represents the overall distribution of transition probabilities across all dispensers. A higher diversity value indicates that transitions are more evenly spread across dispensers, meaning that no single transition overwhelmingly dominates the movement pattern. Conversely, lower diversity suggests that only a few transitions occur more frequently, leading to more repetitive and less varied behavior.

To estimate the variability and compute confidence intervals for global entropy and global transition diversity, we performed a bootstrap resampling procedure with 1,000 iterations. In each iteration, transition probabilities were resampled with replacement, and entropy and diversity values were recomputed. The 95% confidence intervals (CI) were derived from the 2.5th and 97.5th percentiles of the bootstrap distributions.

###### Simulation of Random Behavior for Baseline Comparison

To establish a baseline for global entropy and global transition diversity under random movement conditions, we simulated random sequences of visits and computed their expected entropy and diversity values. A total of 1,000 simulations were performed, each consisting of 1,000 randomly generated sequences of 8 visits (excluding the starting location). In each sequence, dispensers were selected randomly with replacement, ensuring no systematic preference for specific locations. Transition counts were collected across all simulated sequences to compute empirical random transition probabilities. These probabilities were globally normalized to ensure comparability with the observed data. From these simulations, we calculated the mean global entropy and global transition diversity, and derived confidence intervals (at the 95%, 99%, and 99.9% levels) based on the corresponding percentiles of the bootstrap distributions.

###### Permutation Test for Statistical Comparison

Permutation tests were applied to compare the observed global entropy and global transition diversity of each cluster to both the random baseline and between clusters. In 1,000 iterations, the observed data and simulated baseline were pooled, and data were randomly reassigned to two groups of equal size. The mean difference in global entropy and global transition diversity was calculated for each permutation, generating a null distribution. Bonferroni corrections were applied to adjust for multiple comparisons.

###### Transition Matrices

To investigate the behavioral transition patterns, transition matrices were calculated for each cluster, representing the transition probabilities between dispensers across trials. These matrices were computed for both global transitions (including all observed transitions) and significant transitions (those that were significantly different from random chance).

Transition matrices were constructed by counting the frequency of transitions between dispensers across all trials. Transition probabilities were then normalized so that the sum of probabilities for each row (representing outgoing transitions from a dispenser) equaled 1.

To identify statistically significant transitions, binomial tests were conducted for each transition in the matrix, comparing the observed transition probabilities to a uniform random baseline of 1/9 (assuming each dispenser is equally likely to be chosen). Bonferroni correction was applied to adjust for multiple comparisons. Only transitions with corrected p-values less than 0.05 were considered significant.

The similarity between the transition matrices of different clusters was assessed by computing the correlation between their flattened versions. A permutation test with 1,000 iterations was applied to randomly shuffle the values in each matrix pair and compare the correlation of the shuffled matrices to the observed correlation. This process allowed the calculation of a p-value for the correlation, which quantified the likelihood of obtaining the observed correlation by chance. Additionally, the Euclidean distance between the transition matrices was calculated to measure the dissimilarity between matrices, with smaller distances indicating greater similarity.

###### Transition Probability Mapping

To visualize the structure of movement patterns, transition maps were generated for each cluster based on the first 8 choices of each trial. These patterns were visualized using directed graphs where all transition probabilities were represented by a weighted graph, with edge thickness proportional to the transition probability. Significant transitions were represented as a subset of transitions meeting statistical significance.

##### Analysis by Training Phase

###### Transition Probability Computation

The computation of transition probabilities followed the methodology described previously. Briefly, each trial was represented as a sequence of dispenser visits, and transition probabilities were calculated from the frequency of transitions between dispensers. To ensure comparability across training phases, we considered all possible transitions-even those with zero occurrences- and normalized the resulting probabilities on a row-by-row basis (i.e., for each dispenser, the sum of the probabilities of transitions from that dispenser was scaled to equal 1).

###### Global Entropy

Global entropy analysis for training phases was conducted in the same way as described in the previous section. Transition probabilities were globally normalized, so that the sum of all transition probabilities within each cluster equaled 1 and Shannon entropy was computed to measure the unpredictability of transition patterns within each training phase. Bootstrap resampling with 1,000 iterations was used to estimate the variability of entropy values, as described in 6.6.1.2. The 95% confidence intervals (CI) for entropy were derived from the 2.5th and 97.5th percentiles of the bootstrap distributions. Permutation tests were applied to compare the observed entropy between training phases. In 1,000 iterations, data were pooled and randomly reassigned to two groups. The mean difference in entropy was calculated for each permutation, and Bonferroni corrections were applied to adjust for multiple comparisons.

###### Overlap Index Calculation

To assess the degree of similarity between entropy distributions from different training phases, we calculated the Overlap Index. This index measures the shared area between two probability distributions and provides an indication of how similar the transition behaviors are between phases. The index ranges from 0 to 1, with 0 indicating no similarity and 1 indicating complete similarity in the transition patterns. For each pair of training phases, the entropy distributions were calculated, and histograms were constructed with 100 bins, covering the range of observed entropy values. The Overlap Index was then determined by summing, across all bins, the minimum value between the corresponding bins of the two histograms. Higher Overlap Index values indicate greater similarity in the transition patterns between dispensers across training phases

###### Correlation between global entropy and performance rate

To evaluate the linear relationship between the mean performance rate (across the first 8 choices) and the mean global entropy, their correlation was assessed using Pearson’s correlation coefficient across all training phases. Bonferroni correction was applied to account for multiple comparisons. Additionally, a linear regression model was fitted to the data to visually represent and further assess the relationship between these two variables.

###### Transition Matrices

Transition matrices for each training phase were calculated following the same approach as described in the previous section. Significant transitions were identified using binomial tests, with Bonferroni corrections for multiple comparisons.

###### Transition Probability Mapping

Transition maps were generated for each training phase in the same manner as described in the previous section. Significant transitions were identified based on binomial tests, and Bonferroni corrections were applied for multiple comparisons.

###### Local Entropy

Local entropy quantifies the unpredictability of transitions at the individual dispenser level. For each dispenser, we first calculated the transition probabilities, that is, the likelihood of transitioning from that dispenser to each of the other dispensers. The local entropy for a dispenser was then computed using Shannon’s entropy formula:

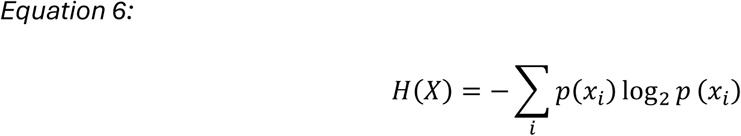

Where *p*(*x_i_*) represents the probability of transitioning to dispenser *i*. For dispensers with no outgoing transitions, the entropy was set to zero. In this context, higher local entropy values indicate greater variability (i.e., less predictable and more diverse transitions), whereas lower values suggest more structured, predictable transition patterns.

To assess whether local entropy values differed significantly across training phases, we applied the Kruskal-Wallis test. When significant differences were detected, pairwise comparisons were performed using Dunn’s post-hoc test with Bonferroni correction.

In addition, to evaluate whether local entropy values differed from what would be expected under random behavior, we performed a bootstrap resampling analysis at the local level. For each dispenser, we resampled the observed transitions 1,000 times with replacement and computed a distribution of local entropy values. From these distributions, we derived 95% confidence intervals (CI) based on the 2.5th and 97.5th percentiles. This allowed us to determine whether the observed local entropy for each dispenser significantly deviated from chance-level expectations.

###### Error Probability

For each dispenser, we calculated error probability as the proportion of errors that occurred following a visit to that dispenser. Specifically, for each training phase, we counted the number of errors that immediately followed visits to each dispenser and divided that number by the total number of trials in that phase. This provided an estimate of the likelihood of making an error after visiting a given dispenser, relative to the overall number of trials-in other words, the probability of an error in any given trial when that dispenser is involved. To assess whether error probabilities differed significantly across training phases, we applied the Kruskal-Wallis test. When significant differences were detected, Dunn’s post-hoc test with Bonferroni correction was used to identify which specific phases differed.

###### Correlation between Local Entropy and Error Probability

To evaluate the linear relationship between mean local entropy and mean error probability, Pearson’s correlation coefficient was computed using data from all training phases combined. A Bonferroni correction was applied to adjust for multiple comparisons. Additionally, a linear regression model was fitted to the data to visually represent and further assess the relationship between these two variables.

#### Markov Model Analysis

The Markov model was trained exclusively on error-free trials to focus on transition patterns observed during successful task performance. This section describes the methods used to construct the transition matrix, predict transitions, evaluate performance, and compare the Markov model to a random baseline.

##### Transition Prediction and Accuracy

Each trial was represented as a sequence of visits to dispensers. Transition probabilities between dispensers were computed by tracking the frequency of transitions from one dispenser to another across all trials. The transition probability from dispenser *i* to dispenser *j* was defined following *Equation 3*.

After calculating these transition frequencies, each row of the transition matrix was normalized to ensure that the sum of outgoing transition probabilities from each dispenser was equal to 1. This normalization ensures that the transition matrix represents valid probability distributions for the Markov model, where each row is a probability distribution over the next possible dispensers.

###### Transition prediction Procedure

To evaluate the model’s predictive capabilities, we compared the Markov model against a random baseline model. For the Markov model, the next dispenser choice was predicted based on the transition matrix computed from the data. Specifically, the prediction of the next dispenser for a given state *i* (current dispenser) is made by selecting the dispenser *j* with the highest transition probability:

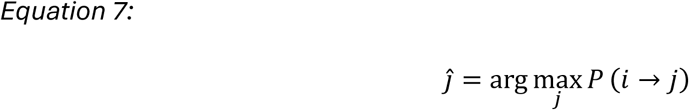

Where *Ĵ* is the predicted next dispenser and *P*(*i* → *j*) is the transition probability from dispenser *i* to dispenser *j*. As a baseline, a random model was implemented, where the next dispenser was selected uniformly at random from all dispensers. Prediction accuracy was calculated as the proportion of correct predictions, i.e., the proportion of times the predicted next dispenser matched the actual next dispenser.

###### Cross-Validation

To evaluate the robustness of the Markov model, we performed 10-fold cross-validation. The dataset was randomly split into 10 subsets (folds), with 9 folds used for training and the remaining fold used for testing. This process was repeated 10 times, with each fold serving as the test set once. For each fold, the transition matrix was computed based on the training data, and the prediction accuracy for both the Markov model and a random baseline model was evaluated on the test set. Performance metrics, including prediction accuracy, were then averaged across all folds to provide a reliable measure of the model’s generalization capability.

In addition, confusion matrices were generated for each fold to analyze the predictions made by the Markov model. The final confusion matrix was obtained by averaging the confusion matrices across all folds, and each row was normalized so that the sum of probabilities for transitions from each actual dispenser equals 1. This normalization accounts for variations in transition frequencies and provides a clear picture of the model’s performance on a per-dispenser basis.

Similarly, an error matrix was constructed by aggregating the counts of mispredicted dispensers across all folds. The error matrix was then normalized globally-by dividing each entry by the total number of errors across all folds-to yield a measure of the relative frequency of errors associated with each dispenser. The normalized error matrix highlights which dispensers are most prone to errors, offering insights into specific weaknesses in the model’s predictions.

###### Evaluation and Statistical Comparison Between Models

To determine whether the Markov model significantly outperformed the random baseline in predicting individual transitions, we conducted a Wilcoxon signed-rank test on the accuracy scores obtained across the cross-validation folds. For both models, mean transition prediction accuracy was computed across folds to provide an overall performance measure. Variability was quantified by calculating the standard deviation and 95% confidence intervals (CI) of these accuracy scores. The Wilcoxon signed-rank test was then applied to assess whether the Markov model’s transition accuracy was significantly higher than that of the random model, ensuring a robust statistical comparison.

###### Transition Matrix Visualization

To visualize the transition patterns learned by the Markov model, we generated predicted transition maps. These maps were represented as directed graphs, where nodes corresponded to dispensers and edges depicted the predicted transitions. The thickness of each edge was proportional to the corresponding transition probability, providing an intuitive visual representation of the local behavioral patterns captured by the model.

###### Sequence Prediction and Accuracy

To further evaluate the performance of the Markov model, we assessed its ability to predict full sequences of dispenser visits. Specifically, we focused on predicting sequences of 8 consecutive choices made by the marmoset. Instead of predicting transitions independently, we treated each sequence as a complete set of transitions to be predicted. Importantly, the same transition matrix computed from the training data in Section 6.7.1 was used for sequence predictions, ensuring consistency across both levels of analysis.

###### Sequence Prediction Procedure

For each sequence, predictions were generated using the same transition matrix from the training phase. The sequence was predicted iteratively, beginning with the first dispenser and selecting the next dispenser at each step based on the highest transition probability. Sequence prediction accuracy was then calculated as the proportion of correctly predicted sequences, representing the percentage of cases where the predicted sequence matched the actual sequence across all eight transitions. This approach assessed the Markov model’s ability to capture global dependencies between transitions. To establish a baseline for comparison, a random model was also evaluated, where each dispenser was predicted with equal probability across all possible dispensers. The accuracy of the random model was computed using the same methodology, allowing for a direct performance comparison.

###### Cross-Validation for Sequence Prediction

To evaluate the generalization ability of the Markov model for sequence prediction, we applied 10-fold cross-validation, ensuring consistency with the methodology used in the previous sections. The dataset was randomly divided into 10 folds, with each fold serving as a test set once while the remaining folds were used for training. For each fold, the transition matrix was computed from the training sequences and used to predict sequences in the test set. Prediction accuracy was assessed for both the Markov and random models, allowing for a comparative evaluation of performance.

###### Evaluation and Statistical Comparison Between Models

To determine whether the Markov model significantly outperformed the random baseline in predicting sequences of dispenser visits, we conducted a Wilcoxon signed-rank test on sequence accuracy scores. Mean sequence accuracy across folds was computed for both models, providing an overall measure of prediction performance. Variability was quantified by calculating the standard deviation and 95% confidence intervals (CI) of these accuracy scores. The Wilcoxon signed-rank test was then applied to assess whether the Markov model’s sequence accuracy was significantly higher than that of the random model, ensuring a robust statistical comparison.

###### Transition Matrix Visualization

To illustrate the predicted sequences, we constructed directed graphs in which nodes represent dispensers and edges represent the corresponding transition probabilities. In these graphs, only the sequence predicted by the Markov model was displayed. The thickness of each edge was scaled according to its transition probability, thereby highlighting the most likely transitions. This visualization provides an intuitive depiction of the predicted sequence and of how effectively the model captures global behavioral patterns.

##### Transition Prediction Accuracy Across Training Phases

To assess the evolution of the Markov model’s performance over time, we evaluated its predictive accuracy separately for each training phase. This analysis allowed us to examine how the model adapted across different learning phases and whether prediction accuracy improved as training progressed.

###### Local Transition Prediction and Accuracy by Training Phase

The same Markov modeling approach described in previous sections was applied independently to each training phase, with separate transition matrices computed for each phase. This ensured that any variations in transition dynamics across training could be captured. For each training phase, we calculated transition probabilities, predicted the next dispenser based on the highest transition probability, and compared the Markov model’s accuracy to a random baseline.

###### Cross-Validation by Training Phase

For cross-validation, the number of folds was dynamically adjusted based on the available data in each training phase. When sufficient sequences were available, 10-fold cross-validation was performed, otherwise, the number of folds was reduced accordingly to ensure reliable evaluation. Accuracy scores were averaged across folds to provide a robust measure of generalization within each training phase. Additionally, confusion matrices were computed separately for each training phase to analyze errors and evaluate model performance over time.

###### Evaluation and Statistical Comparison Between Models by Training Phase

To determine whether the Markov model’s performance improved throughout training, we compared accuracy across training phases as well as against the random baseline for each phase. Variability was quantified by calculating the standard deviation and 95% confidence intervals (CI) of these accuracy scores. For each training phase, a Wilcoxon signed-rank test with Bonferroni correction was conducted to assess whether the Markov model significantly outperformed the random model. In addition, a Kruskal-Wallis test was performed to evaluate whether differences in accuracy across training phases were statistically significant. When the Kruskal-Wallis test indicated significant differences, Dunn’s post-hoc test with Bonferroni correction was applied to identify which specific phases differed.

###### Transition Matrix Visualization by Training Phase

To visualize how the Markov model’s predictions evolved over training, we generated predicted transition maps for each training phase. These maps were represented as directed graphs, where nodes corresponded to dispensers and edges depicted the predicted transitions for that phase. The thickness of each edge was proportional to the corresponding transition probability, providing an intuitive visual representation of the transition patterns during each phase. This approach allowed us to track the evolution of local behavioral patterns as training progressed.

##### Sequence Prediction Accuracy Across Training Phases

Beyond individual transitions, we also evaluated the Markov model’s ability to predict full sequences of dispenser visits separately for each training phase. This analysis allowed us to assess whether the model became more effective at capturing global behavioral patterns over time. Importantly, the same transition matrices computed from the training data for each training phase in Section 6.7.3 were used for sequence predictions, ensuring consistency across both levels of analysis.

###### Sequence Prediction and Cross-Validation by Training Phase

For each training phase, sequences were extracted and analyzed separately, following the same Markov modeling approach as in Section 6.7.2. We used the transition matrices computed in Section 6.7.3 for each training phase. This ensured consistency across both transition- and sequence-level analyses.

Cross-validation was adapted dynamically to match the dataset size in each training phase. 10- fold cross-validation was applied when enough sequences were available, while fewer folds were used when necessary to maintain a robust evaluation. In each fold, the precomputed transition matrix from the training phase was used to predict full sequences of up to 8 transitions in the test set. The Markov model’s sequence accuracy was compared to that of a random baseline model, which predicted each transition with equal probability.

###### Evaluation and Statistical Comparison of Sequence Prediction

To assess whether the Markov model’s sequence prediction accuracy significantly improved over training, a Wilcoxon signed-rank test with Bonferroni correction was conducted for each training phase to determine if the Markov model significantly outperformed the random baseline. Additionally, a Kruskal-Wallis test was applied to analyze whether sequence prediction accuracy evolved significantly across training phases. When the Kruskal-Wallis test indicated significant differences, Dunn’s post-hoc test with Bonferroni correction was used to identify which specific training phases differed.

###### Sequence Transition Visualization

To illustrate the predicted sequences for each training phase, we constructed directed graphs in which nodes represent dispensers and edges represent the corresponding transition probabilities. In these graphs, only the sequence predicted by the Markov model was displayed. The thickness of each edge was scaled according to its transition probability, thereby highlighting the most likely transitions for each training phase.

#### Analysis of Sequence Variability and Similarity

To further explore the structure of dispensers transition sequences, we analyzed the variability and similarity of observed sequences across error-free trials.

##### Identification of Unique Sequences and Frequency Analysis

Transition sequences were extracted from error-free trials, where each sequence was defined as a structured set of consecutive transitions between dispensers. The number of unique sequences was identified, and their relative usage frequencies were computed to evaluate the diversity of sequence patterns. To determine whether certain sequences were dominant behavioral patterns, we identified the most frequently used sequence for each animal and calculated its proportion relative to all observed sequences.

##### Analysis of Sequence Diversity

To quantify the diversity of navigation sequences used in error-free trials, we computed the normalized Shannon diversity index, which is derived from Shannon entropy and normalized by its theoretical maximum. The normalized Shannon index was defined as:

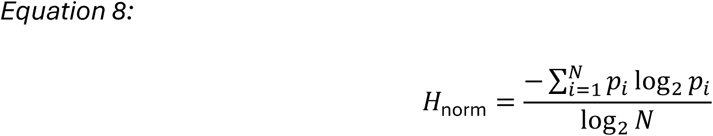

where N is the total number of unique sequences, and *p_i_* represents the relative frequency of sequence *i*, computed as the number of occurrences of that sequence divided by the total number of sequences.

This normalization ensures that *H*_norm_ ranges between 0 (when only one sequence is used, indicating no diversity) and 1 (when all sequences occur with equal probability, indicating maximum diversity).

##### Analysis of Sequence Similarity

To assess the structural similarity between sequences, we examined common transitions, defined as pairs of consecutive dispensers appearing within sequences. Sequence similarity was quantified using the Jaccard similarity index, calculated as:

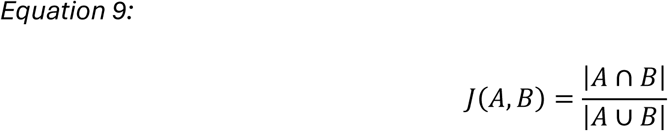

Where A and B represent the sets of transitions in two sequences, and |A ∩ B| denotes the number of shared transitions between them. This index varies between 0 and 1, with 0 indicating no shared transitions and 1 indicating complete overlap. This measure provided insight into the degree of transition overlap, indicating whether sequences followed similar transition patterns or exhibited high variability.

We computed the mean Jaccard similarity across all sequence pairs and analyzed the distribution of similarity levels. Specifically, we determined the proportion of sequence pairs exceeding similarity thresholds ranging from 0.1 to 1.0, providing a detailed view of how frequently sequences shared a given percentage of their transitions. This approach allowed us to evaluate sequence overlap and assess the structural consistency of navigation patterns over trials.

To further examine sequence similarity patterns, the Jaccard similarity matrix was represented as a heatmap, where each cell indicated the similarity score between two sequences. This visualization provided a structured overview of sequence similarity distributions and facilitated the identification of clusters of highly similar sequences.

Additionally, to quantify the occurrence of individual transitions, we counted the frequency of each transition (i.e., movement from one dispenser to another) across all sequences. These frequencies were normalized by computing the percentage of sequences containing each transition relative to the total number of sequences analyzed.

##### Sequence Diversity Across Training Phase

To examine the evolution of sequence diversity over training, we computed the normalized Shannon diversity index separately for each training phase (as described in Section 6.8.2). To model the trajectory of diversity across training, we applied weighted least squares (WLS) quadratic regression, with weights determined by the number of trials in each phase to account for sample size differences. We compared the performance of a simple linear regression to that of the quadratic model to determine whether a non-linear (e.g., U-shaped) trend provided a better fit. Model fit was evaluated using R^2^ values, and the statistical significance of the quadratic term was examined. Prior to interpreting the results, we verified that the model assumptions-normality of residuals, homoscedasticity, independence of errors, and absence of multicollinearity-were met, confirming the reliability of our approach.

##### Dominant Sequence Frequency, New Sequence Ratio, and Reused Sequence Ratio Across Training Phases

To examine how sequence selection evolved throughout training, we quantified three key metrics: dominant sequence frequency, new sequence ratio, and reused sequence ratio. These measures were computed separately for each training phase to assess changes in the patterns of sequence selection over time.

###### Dominant Sequence Frequency

The dominant sequence was identified as the most frequently occurring sequence across all error-free trials. For each training phase, we computed its frequency of use, defined as the proportion of trials in which the dominant sequence was executed.

###### New Sequence Ratio

The New Sequence Ratio (NSR) was computed as the number of unique sequences in each training phase that had not been observed in any previous phase. This metric was calculated iteratively by maintaining a cumulative set of previously encountered unique sequences and identifying, for each phase, the fraction of new unique sequences relative to all unique sequences observed during that phase

###### Reused Sequence Ratio

The Reused Sequence Ratio (RSR) was computed as the proportion of sequences in each training phase that had already been observed in previous phases. For each training phase, reused sequences were identified by comparing the sequences in the current phase to a cumulative set of sequences encountered in earlier phases. The reused sequence ratio was then calculated as the total number of reused sequences divided by the total number of sequences in that phase.

##### Linear Relationship Between Sequence Diversity and Sequence Selection Strategies

To assess the relationship between sequence diversity and sequence selection patterns, we conducted linear regression analyses using the normalized Shannon Diversity Index as the independent variable and either the New Sequence Ratio (NSR) or the Reused Sequence Ratio (RSR) as the dependent variable. Separate ordinary least squares (OLS) regression models were fitted for each metric and each animal to evaluate whether higher diversity was associated with a greater tendency to generate novel sequences (NSR) or reuse existing ones (RSR). Model fit was assessed using R-squared values, and 95% confidence intervals were computed to visualize prediction uncertainty. To validate the models, residual normality was assessed using the Shapiro-Wilk test, independence of errors was verified using the Durbin-Watson statistic, and homoscedasticity was tested using the Breusch-Pagan test.

##### Sequence Similarity Within and Across Training Phases

To assess how sequence similarity evolved throughout training, we applied the same Jaccard similarity analysis described in previous sections computed separately for each training phase. For each phase, sequences were extracted and the pairwise Jaccard similarity index was calculated to quantify the overlap in transitions between sequences. Mean similarity scores and SEM were computed to evaluate whether sequences became more homogeneous or remained diverse over time. Additionally, we assessed sequence similarity between training phases by computing pairwise Jaccard similarity scores between sequences belonging to different phases. This analysis allowed us to examine whether similar sequences tended to reoccur in later phases, indicating consolidation or reuse of specific patterns. For visualization, Jaccard similarity matrices were generated for each phase and across-phase comparisons, and represented as heatmaps to illustrate the structural similarity of transition patterns over time. To determine whether sequence similarity differed significantly across phases, Kruskal-Wallis tests were performed on similarity scores, when significant effects were found, Dunn’s post-hoc tests with Bonferroni correction were applied to identify specific pairwise differences.

