## Supplementary for "Structured navigation emerges from self-guided spatial learning in freely moving common marmosets"

### SUPPLEMENTARY MATERIALS

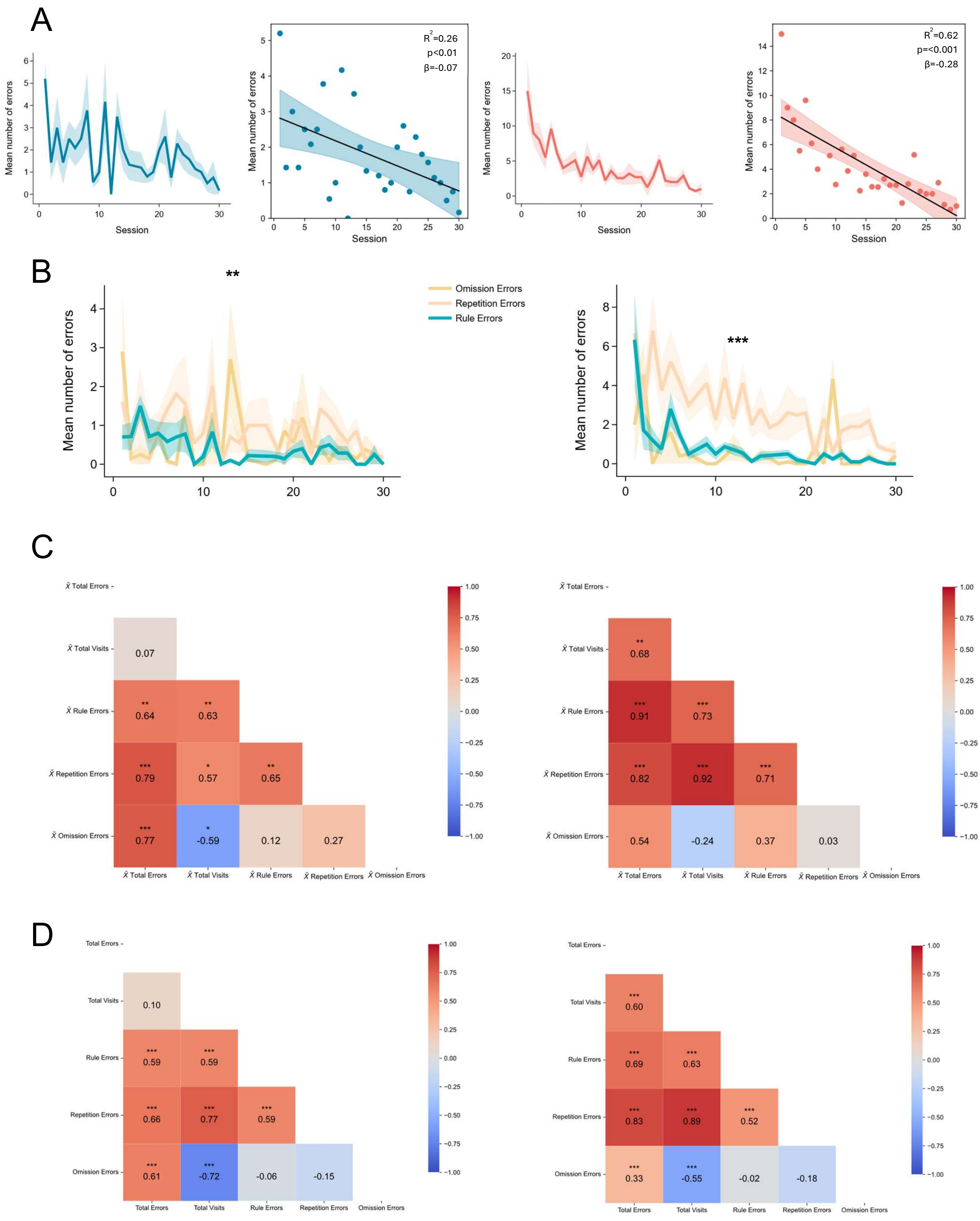
A
Mean number of errors
Session
R2=0.26
p<0.01
β=-0.07
R2=0.62
p=<0.001
β=-0.28
B
**
***
Mean number of errors
Omission Errors
Repetition Errors
Rule Errors
C
X̄ Total Errors
X̄ Total Visits
X̄ Rule Errors
X̄ Repetition Errors
X̄ Omission Errors
0.07
0.64
0.63
0.79
0.57
0.65
0.77
-0.59
0.12
0.27
0.68
0.91
0.73
0.82
0.92
0.71
0.54
-0.24
0.37
0.03
D
Total Errors
Total Visits
Rule Errors
Repetition Errors
Omission Errors
0.10
0.59
0.59
0.66
0.77
0.59
0.61
-0.72
-0.06
-0.15
0.60
0.69
0.63
0.83
0.89
0.52
0.33
-0.55
-0.02
-0.18

**Figure S1. Marmoset performance improves over sessions, while error types follow distinct dynamics**

**A. Mean number of errors across training sessions.** *Left*: line plots showing the session-wise mean number of errors for Animal E (blue) and Animal F (orange), with shaded areas representing SEM. *Right*: corresponding linear regression analyses of error rates across sessions, with dots indicating session means, black lines showing best-fit regressions, and shaded areas the 95% confidence intervals.

**B. Evolution of error types over sessions**. Line plots of session-wise means for rule, repetition, and omission errors for each animal. Shaded areas indicate SEM. Statistical analyses were performed using the Kruskal-Wallis test followed by Dunn's post-hoc test with Bonferroni correction. Significant p-values from the Kruskal-Wallis test are marked with asterisks (** $p < 0.01$, ***$p < 0.001$).

**C. Session-level correlation matrix.** Pearson correlations between session-wise means of rule, repetition, and omission errors, total visits, and total errors. Color intensity represents correlation strength (-1 to +1). Significant correlations after Bonferroni correction are marked with asterisks*** $p < 0.001$, ** $p < 0.01$.

**D. Trial-level correlation matrix.** Same as in C, but computed at the trial level. Color scale and significance markers as above.

| | ANIMAL E | | | ANIMAL F | | |
|---|---|---|---|---|---|---|
| **Variable** | **Coefficient** | **Std Error** | **p-value (Bonferroni)** | **Coefficient** | **Std Error** | **p-value (Bonferroni)** |
| #days since last training session | 0.02415281 | 0.06072828 | 1 | 0.05822239 | 0.09103679 | 1 |
| experimenter_encoded | -0.06338979 | 0.37408384 | 1 | -0.76446322 | 0.80390622 | 1 |
| **session_numeric** | **-0.06797315** | **0.0239631** | **0.03647502** | **-0.2589218** | **0.03201119** | **1.04E-07** |
| num_trials | 0.02970938 | 0.11353353 | 1 | -0.10058889 | 0.2747594 | 1 |

**Table S1: Multiple linear regression assessing factors influencing task performance**

Summary of a multiple linear regression model predicting the mean number of total errors per session. The table reports coefficient estimates, robust standard errors (HC3), and Bonferroni-corrected p-values for each predictor: number of days since the last session, experimenter identity (encoded), session number, and number of trials. The model accounted for 27% of the variance in performance for Animal E ($R^2 = 0.27$) and 64% for Animal F ($R^2 = 0.64$). After correction, only session number had a significant effect (Animal E: $\beta = -0.07$, $p = 0.02$; Animal F: $\beta = -0.26$, $p < 0.001$), while all other variables showed no significant contribution ($p > 0.05$). These results indicate that performance improvements were primarily driven by training progression, independent of other contextual factors.

A

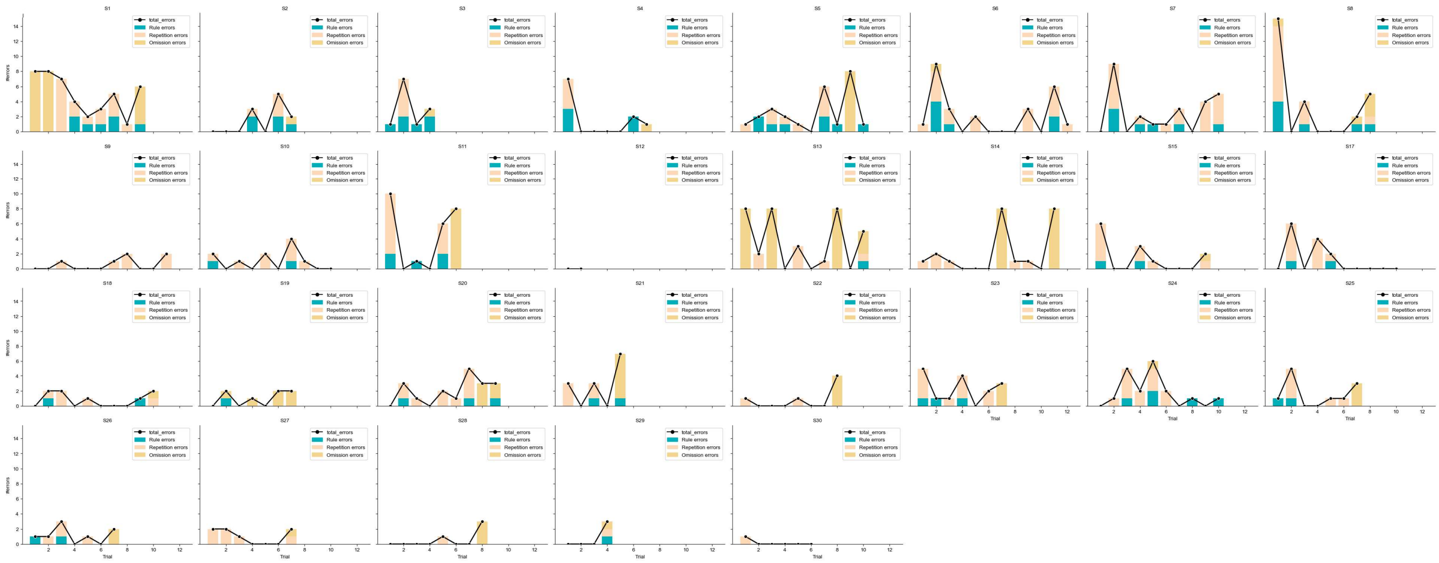


B

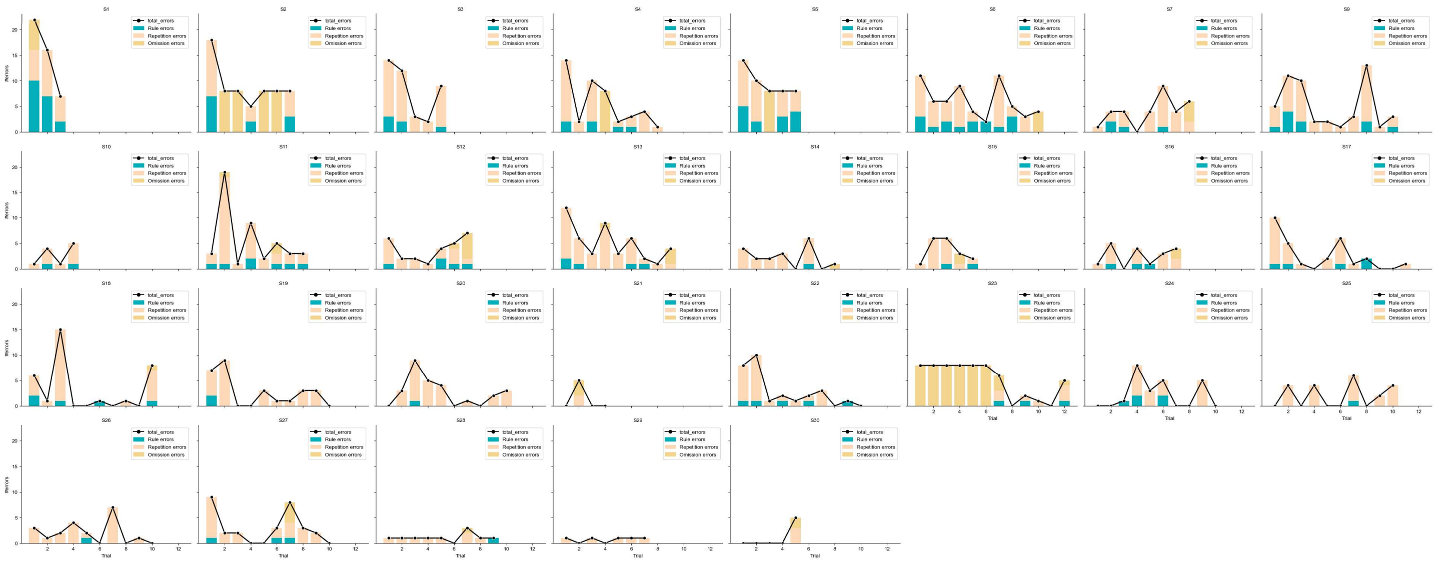


**Fig. S2. Error type dynamics across trials and sessions**
Stacked bar plots showing the number and type of errors per trial across sessions (each facet represents one session). Within each trial, error types are color-coded and stacked: rule errors (blue), repetition errors (beige), and omission errors (yellow). The total number of errors per trial is overlaid as a black line with markers. Data are shown for Animal E (panel A) and Animal F (panel B).

| | ANIMAL E | | ANIMAL F | |
|---|---|---|---|---|
| cluster | omission_errors | total_nb_visits | omission_errors | total_nb_visits |
| 0 | 0.06 ± 0.02 | 8.49 ± 0.06 | 0.02 ± 0.01 | 9.65 ± 0.12 |
| 1 | 0.08 ± 0.04 | 12.97 ± 0.36 | 0.07 ± 0.04 | 17.37 ± 0.48 |
| 2 | 7.8 ± 0.2 | 0.25 ± 0.25 | 8.0 ± 0.0 | 0.0 ± 0.0 |
| 3 | 2.92 ± 0.26 | 5.58 ± 0.31 | 3.33 ± 0.38 | 7.91 ± 1.02 |

**Table S2: Mean values of clusters centroids.**
Table reporting the mean (± SEM) coordinates of cluster centroids identified through K-means clustering based on two metrics: omission errors and total number of visits. Each trial was assigned to one of four clusters, and the centroid values are shown for each cluster and for both Animal E and Animal F.

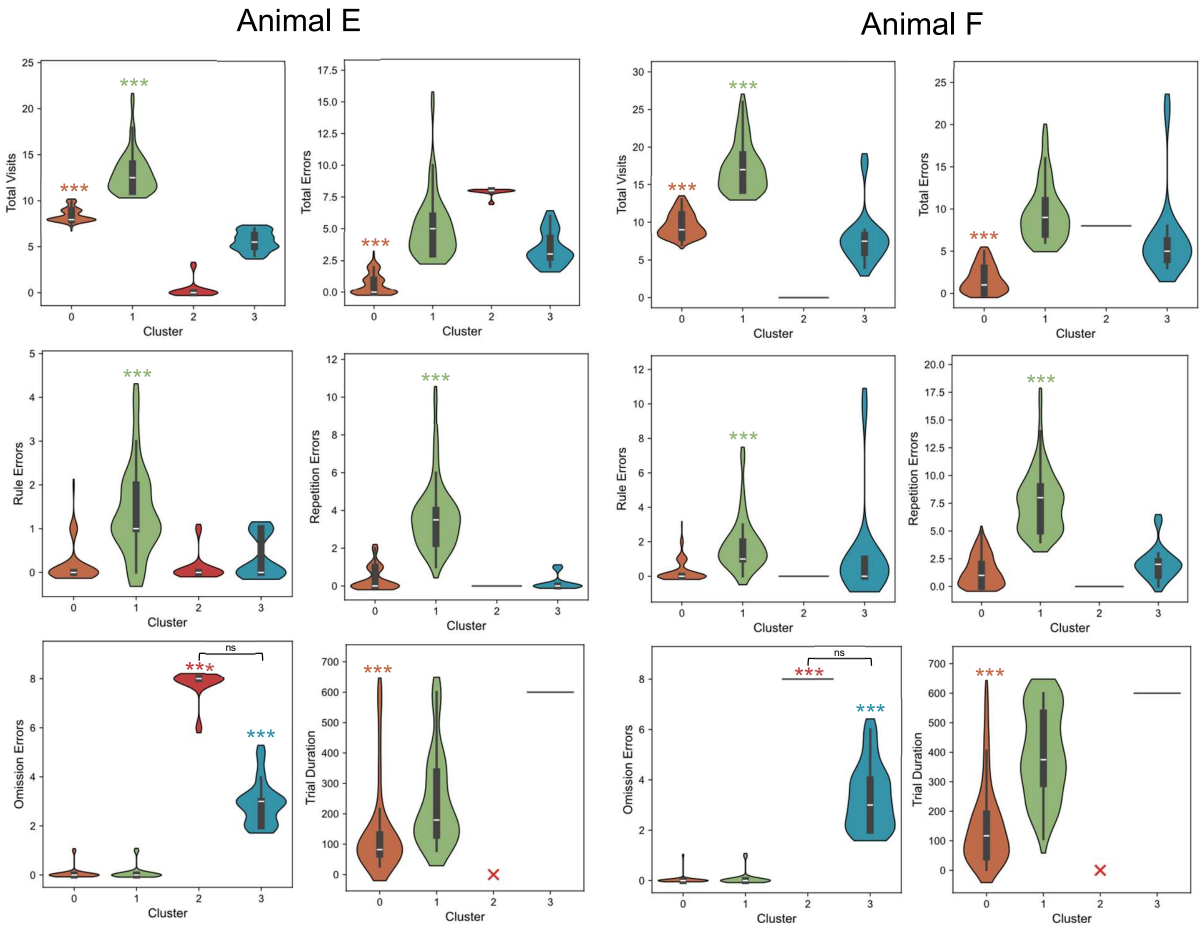


**Fig. S3: Behavioral differences across clusters.**
Violin plots showing the distribution of key task performance metrics across the four clusters identified by K-means clustering. Each panel represents a different variable (total number of visits, total errors, rule errors, repetition errors, omission errors, and trial duration) illustrating the data spread and median within each cluster. The width of each violin indicates the density of observations, and the horizontal bar marks the median. Statistical differences across clusters were assessed using the Kruskal–Wallis test, followed by Dunn's post hoc comparisons with Bonferroni correction. Significant differences are indicated with asterisks (*** $p < 0.001$). Red crosses denote clusters for which no data were available.

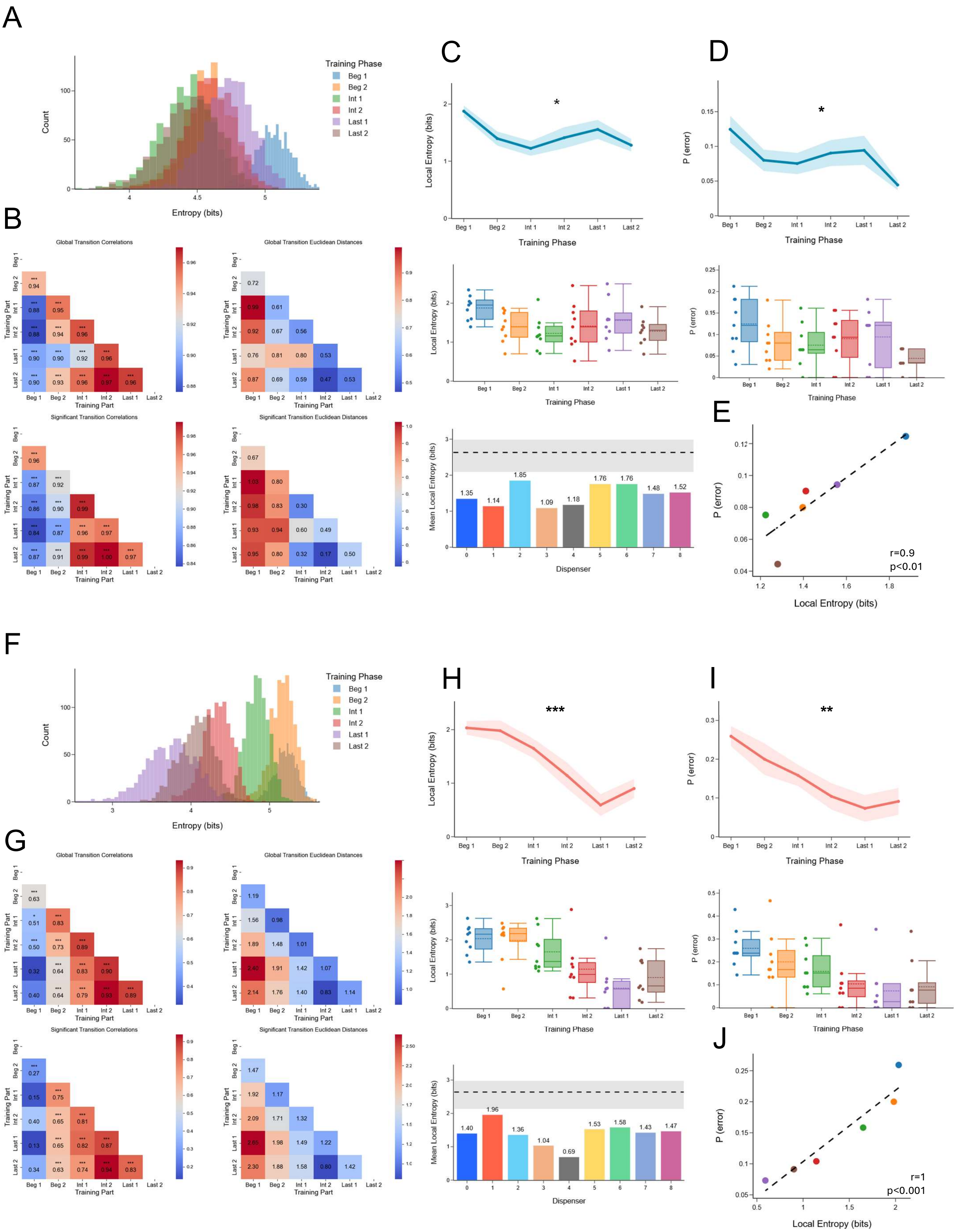
A
Training Phase
Beg 1
Beg 2
Int 1
Int 2
Last 1
Last 2
Count
Entropy (bits)
B
Global Transition Correlations
Global Transition Euclidean Distances
Significant Transition Correlations
Significant Transition Euclidean Distances
Training Part
C
Local Entropy (bits)
Training Phase
D
P (error)
Training Phase
Mean Local Entropy (bits)
Dispenser
E
P (error)
Local Entropy (bits)
r=0.9
p<0.01
F
Training Phase
Count
Entropy (bits)
G
Global Transition Correlations
Global Transition Euclidean Distances
Significant Transition Correlations
Significant Transition Euclidean Distances
Training Part
H
Local Entropy (bits)
Training Phase
I
P (error)
Training Phase
Mean Local Entropy (bits)
Dispenser
J
P (error)
Local Entropy (bits)
r=1
p<0.001

**Fig. S4: Evolution of entropy and transition structure across training phases.**
*(A–E) Analyses for Animal E; (F–J) Corresponding analyses for Animal F.*
**A, F. Distribution of global entropy across training phases.** Overlaid histograms showing the bootstrapped entropy distributions of error-free trials for each training phase. Color-coded curves indicate progressive shifts in entropy across training.
**B, G. Global and significant transition pattern similarity.**
**Top**. *Leff*: Heatmaps showing Pearson correlations between global transition matrices across training phases. Color intensity reflects the strength of the correlation, with asterisks indicating statistically significant values (Bonferroni-corrected). *Right*: Euclidean distance matrices between global transition patterns, with darker shades indicating greater dissimilarity between phases.
**Bottom**: Same analyses restricted to significant transitions only, i.e., transitions occurring above chance ($p = 1/9$; binomial test with Bonferroni correction). *Left*: Correlation matrices of significant transitions; *Right*: Euclidean distances between phases based on significant transitions.
**C, H. Local entropy dynamics across training.**
**Top:** Line plots showing the evolution of mean local entropy (± SEM) across training phases. Asterisks indicate significant differences across phases (Kruskal–Wallis test; * $p < 0.05$, *** $p < 0.001$).
**Middle:** Box plot showing the distribution of local entropy (in bits) across the six training phases. Boxes represent the interquartile range (IQR), medians are shown as horizontal lines, and whiskers extend to 1.5× IQR. Individual data points are shown as dots with horizontal jitter for better visibility.
**Bottom:** Mean local entropy per dispenser, averaged across sessions. Colored bars show observed values; dashed lines represent the entropy expected under random transitions, with shaded areas indicating the 95% confidence interval of this baseline.
**D, I. Error probability across training.**
**Top**: Line plots showing the evolution of mean error probability (± SEM) across training phases, computed as the likelihood of making an error after visiting a dispenser. Asterisks indicate significant differences between phases (Kruskal–Wallis test; * $p < 0.05$, ** $p < 0.01$).
**Bottom**: Boxplots of error probability per phase, with the same conventions as above.
**E, J. Correlation between entropy and error probability.**
Scatterplots showing the relationship between mean local entropy and mean error probability across training phases. Each dot represents one phase, color-coded accordingly. Dashed lines indicate the linear regression fit.

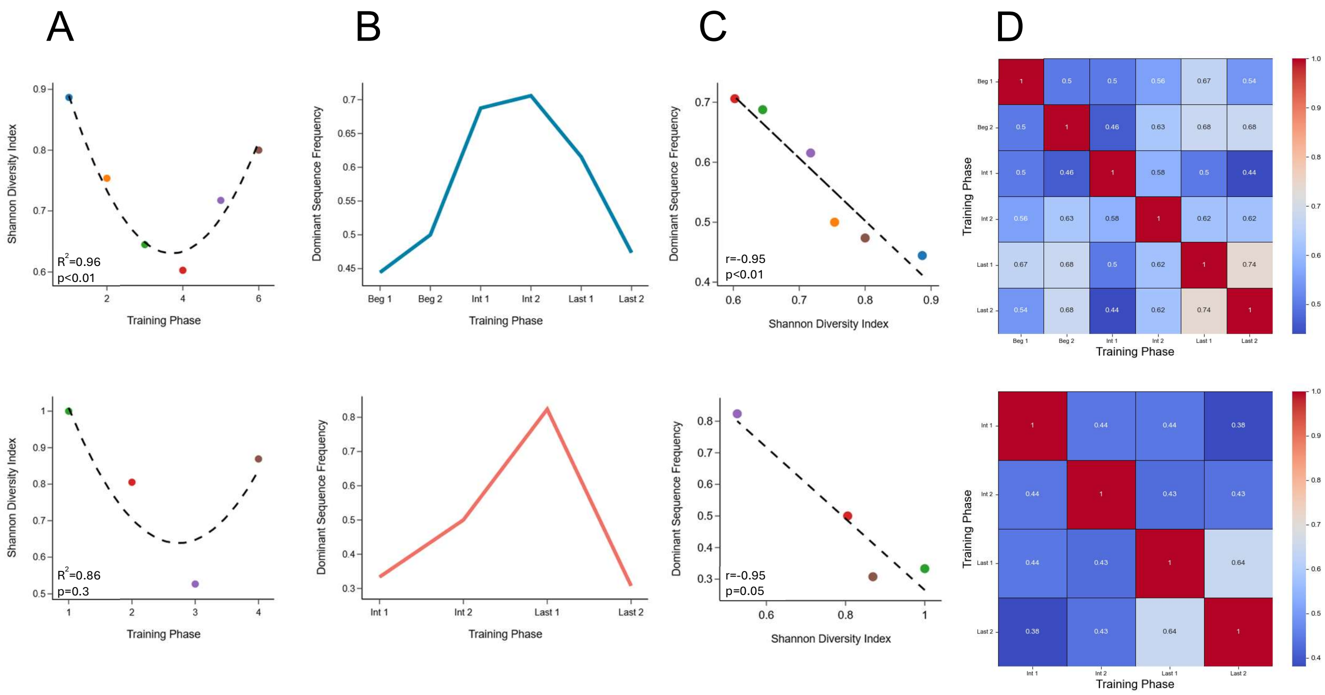


**Fig. S5: Dynamics of sequence diversity, dominance, and structural similarity across training**

**A. Quadratic fit of sequence diversity across training phases.** Scatter plots showing the normalized Shannon diversity index across training phases. Each point represents a phase and is color-coded accordingly. Dashed lines indicate the best-fit quadratic regression models, illustrating the non-linear trajectory of sequence diversity over training. The x-axis reflects the six ordered training phases (Beg 1 to Last 2 for Animal E; Int 1 to Last 2 for Animal F). $R^2$ and p-values indicate the fit quality of the model. *Top*: Animal E; *Bottom*: Animal F.

**B. Frequency of the dominant sequence across training.** Line plots showing the proportion of error-free trials in which the most frequently used sequence appeared during each training phase. Increases indicate growing reliance on a specific behavioral motif. *Top*: Animal E; *Bottom*: Animal F.

**C. Correlation between dominant sequence frequency and sequence diversity.** Scatter plot illustrating the relationship between the Shannon diversity index and the frequency of the dominant sequence across training phases. Points are color-coded by training phase. The dashed regression line highlights the trend, with the Pearson correlation coefficient (r) and p-value reporting the statistical association. *Top*: Animal E; *Bottom*: Animal F.

**D. Similarity of sequence composition between training phases.** Heatmaps representing the Jaccard similarity index between the sets of unique error-free sequences observed in each training phase. Higher values (warmer colors) indicate greater overlap in sequence composition, revealing the temporal consolidation of dominant patterns. *Top*: Animal E; *Bottom*: Animal F.
